# ATP disrupts lipid binding equilibrium to drive retrograde transport critical for bacterial outer membrane asymmetry

**DOI:** 10.1101/2021.05.25.445566

**Authors:** Wen-Yi Low, Shuhua Thong, Shu-Sin Chng

## Abstract

The hallmark of the Gram-negative bacterial envelope is the presence of the outer membrane (OM). The OM is asymmetric, comprising lipopolysaccharides (LPS) in the outer leaflet and phospholipids (PLs) in the inner leaflet; this critical feature confers permeability barrier function against external insults, including antibiotics. To maintain OM lipid asymmetry, the OmpC-Mla system is believed to remove aberrantly localized PLs from the OM and transport them to the inner membrane (IM). Key to the system in driving lipid trafficking is the MlaFEDB ABC transporter complex in the IM, but mechanistic details, including transport directionality, remain enigmatic. Here, we develop a sensitive point-to-point in vitro lipid transfer assay that allows direct tracking of [^14^C]-labelled PLs between the periplasmic chaperone MlaC and MlaFEDB reconstituted into nanodiscs. We reveal that MlaC spontaneously transfers PLs to the IM transporter in an MlaD-dependent manner that can be further enhanced by coupled ATP hydrolysis. In addition, we show that MlaD is important for modulating productive coupling between ATP hydrolysis and such retrograde PL transfer. We further demonstrate that spontaneous PL transfer also occurs from MlaFEDB to MlaC, but such anterograde movement is instead abolished by ATP hydrolysis. Our work uncovers a model where PLs reversibly partition between two lipid binding sites in MlaC and MlaFEDB, and ATP binding and/or hydrolysis shift this equilibrium to ultimately drive retrograde PL transport by the OmpC-Mla system. These mechanistic insights will inform future efforts towards discovering new antibiotics against Gram-negative pathogens.

**Significance Statement:** Biological membranes define cellular boundaries, allow compartmentalization, and represent a prerequisite for life. In Gram-negative bacteria, the outer membrane (OM) prevents entry of toxic substances, conferring intrinsic resistance against many antibiotics. This barrier function requires unequal distribution of lipids across the OM bilayer, yet how such lipid asymmetry is maintained is not well understood. In this study, we established the directionality of lipid transport for a conserved membrane protein complex, and uncovered mechanistic insights into how ATP powers such transport from the OM to the inner membrane. Our work provides fundamental understanding of lipid trafficking within the Gram-negative double-membrane envelope in the context of OM lipid asymmetry, and highlights the importance of targeting lipid transport processes for future antibiotics development.

## Introduction

The cell envelope of Gram-negative bacteria is composed of two lipid bilayers. The inner membrane (IM) is a phospholipid (PL) bilayer while the outer membrane (OM) contains both PLs and lipopolysaccharides (LPS). The OM is asymmetric with LPS residing in the outer leaflet and PLs in the inner leaflet; such lipid asymmetry allows the membrane to function as an effective barrier against the entry of toxic substances, including antibiotics (1, 2). Extensive efforts have contributed towards an improved understanding of the molecular mechanisms involved in OM assembly and homeostasis. We now know that the Lol, Bam, and Lpt pathways are responsible for the unidirectional transport and/or assembly of OM lipoproteins, β-barrel proteins and LPS, respectively (3-5). However, mechanisms for PL transport (6), which occur in both directions for assembly and homeostasis of the OM (7-9), remain largely elusive.

The first pathway implicated in bacterial PL transport is the OmpC-Mla system (10, 11). Cells lacking this pathway accumulate PLs in the outer leaflet of the OM, indicating a role in the maintenance of OM lipid asymmetry. The OmpC-Mla system is believed to mediate retrograde PL transport where the periplasmic MlaC chaperone shuttles PLs from the OmpC-MlaA complex in the OM to the MlaFEDB ATP-binding cassette (ABC) transporter in the IM. MlaA is an OM lipoprotein that interacts and works with trimeric osmoporin OmpC (11, 12), but forms a separate hydrophilic channel in the OM for trans-bilayer PL translocation (12, 13). While MlaA also interacts with OmpF, only removing OmpC, but not OmpF, results in OM lipid asymmetry defects in cells (11); therefore, OmpC is functionally important in the Mla system, but its exact role is unclear. MlaC is found to bind PLs with high affinity in a deep hydrophobic pocket (14-16), presumably enabling extraction of PLs from the OM via OmpC-MlaA. In the IM ABC transporter, MlaE and MlaF constitute the permease and nucleotide binding domains, respectively. MlaD is a substrate-binding protein containing a single transmembrane helix, and its periplasmic MCE domain (17) forms a hexamer with a hydrophobic pore and co-purifies with PLs (14, 18). Recent structural studies of the MlaFEDB complex revealed that the MlaE permease domains complement the MlaD hydrophobic pore to form a contiguous cavity for binding and transporting PLs (19-23). MlaB is a cytoplasmic protein containing the STAS domain (24), and has been shown to be essential for the proper assembly and activity of the IM complex (18, 25). How ATP hydrolysis may drive PL movement within the MlaFEDB complex is not known.

Apart from the lack of mechanistic details, there is still substantial controversy with regards to the directionality of lipid transport mediated by the OmpC-Mla system. Genetic studies in *Escherichia coli* and *Acinetobacter baumannii* are more consistent with the retrograde (OM-to-IM) model (10, 26). In particular, *E. coli mla* or *ompC* mutants accumulate outer leaflet PLs in the OM, a phenotype that can be corrected by overexpression of PldA, the OM phospholipase (10, 11). In addition, *A. baumannii* strains lacking lipooligosaccharides (LOS) have enhanced growth and restored OM barrier function when the Mla system and PldA are removed, presumably because these cells need enough PLs to maintain the outer leaflet of the OM (26, 27). Nevertheless, these experiments in cells lack evidence that demonstrate the direct transport of PLs. While there was a report suggesting the system in *A. baumannii* may be involved in anterograde (IM-to-OM) PL transport (28), it has recently been shown that the *mla* mutant strains used had secondary mutations in them, thus rendering the claim invalid (29). A key experiment that sheds light on transport is when the overexpression of MlaFEDB together with MlaC partially rescued defects in retrograde transport of PLs, observed in *E. coli* strains lacking the Tol-Pal complex (30). Despite this, however, in vitro reconstitution demonstrated ATP-independent (anterograde) transfer of PLs from the MlaFEDB complex (in liposomes) to MlaC (31). These conflicting data throw into question the biological significance of ATP hydrolysis and argue for the need to provide additional in vitro evidence to truly define the directionality of lipid transport mediated by the OmpC-Mla system.

In this study, we utilize [^14^C]-labelled lipids to directly track the point-to-point transfer of PLs between MlaC and nanodisc-reconstituted MlaFEDB complex. We show that PLs bound to MlaC can be spontaneously transferred to MlaFEDB in an MlaD-dependent manner. The presence of ATP further enhances such retrograde transfer; we demonstrate that ATP hydrolysis is coupled to PL transport, and this process too is modulated by MlaD. Interestingly, spontaneous PL transfer can also happen from MlaFEDB to MlaC, but ATP hydrolysis prevents this ‘anterograde’ transfer, ultimately driving PL transport in the retrograde direction. Our work establishes that the OmpC-Mla system powers the retrograde transport of PLs via ATP hydrolysis to maintain OM lipid asymmetry.

## Results

### MlaD stimulates the ATP hydrolytic activity of MlaFEB in a native lipid environment

To gain insights into the directionality of PL transport mediated by the OmpC-Mla system, we focused on lipid transfer between MlaC and the MlaFEDB complex embedded in a lipid bilayer. We first reconstituted MlaFEDB and MlaFEB complexes into lipid nanodiscs comprising *E. coli* polar lipids stabilized by the membrane scaffold protein MSP1E3D1 (Figure 1A) (32), and tested their functionality in ATP hydrolysis. At room temperature, the ATPase activity of nanodisc-embedded MlaFEDB was significantly lower compared to that of the same complex solubilized in detergent micelles (Figure S1A, *k*_cat_: 0.029 ± 0.002 vs 0.126 ± 0.008 μmol ATP s^-1^/μmol complex, respectively), likely due to higher conformational restriction within the membrane. This effect is similar to that observed in MlaFEDB-containing proteoliposomes composed also of native *E. coli* polar lipids (31), but opposite in complexes reconstituted in nanodiscs (21) or proteoliposomes (20) comprising synthetic lipids, suggesting an influence of lipid composition on ATPase function. At physiological temperature (37°C), however, ATP hydrolytic activity was largely recovered in the nanodisc-embedded MlaFEDB complex (*k*_cat_: 0.090 ± 0.002 μmol ATP s^-1^/μmol complex). In contrast to complexes in detergent micelles (18), we further showed that the ATPase activity of nanodisc-embedded MlaFEDB was much higher than that of MlaFEB (*k*_cat_: 0.011 ± 0.002 μmol ATP s^-1^/μmol complex) at 37°C (Figure 1B). In addition, this activity remained unaffected upon addition of holo- (PL-bound) or apo- (PL-free) MlaC (Figure S1B). The MlaFEB structure does not change significantly in the absence of MlaD (22); therefore, the presence of MlaD actually stimulates the ATP hydrolytic function of MlaFEB, consistent with MlaD being the substrate binding protein in direct contact with the ABC transporter.

**Figure 1.**
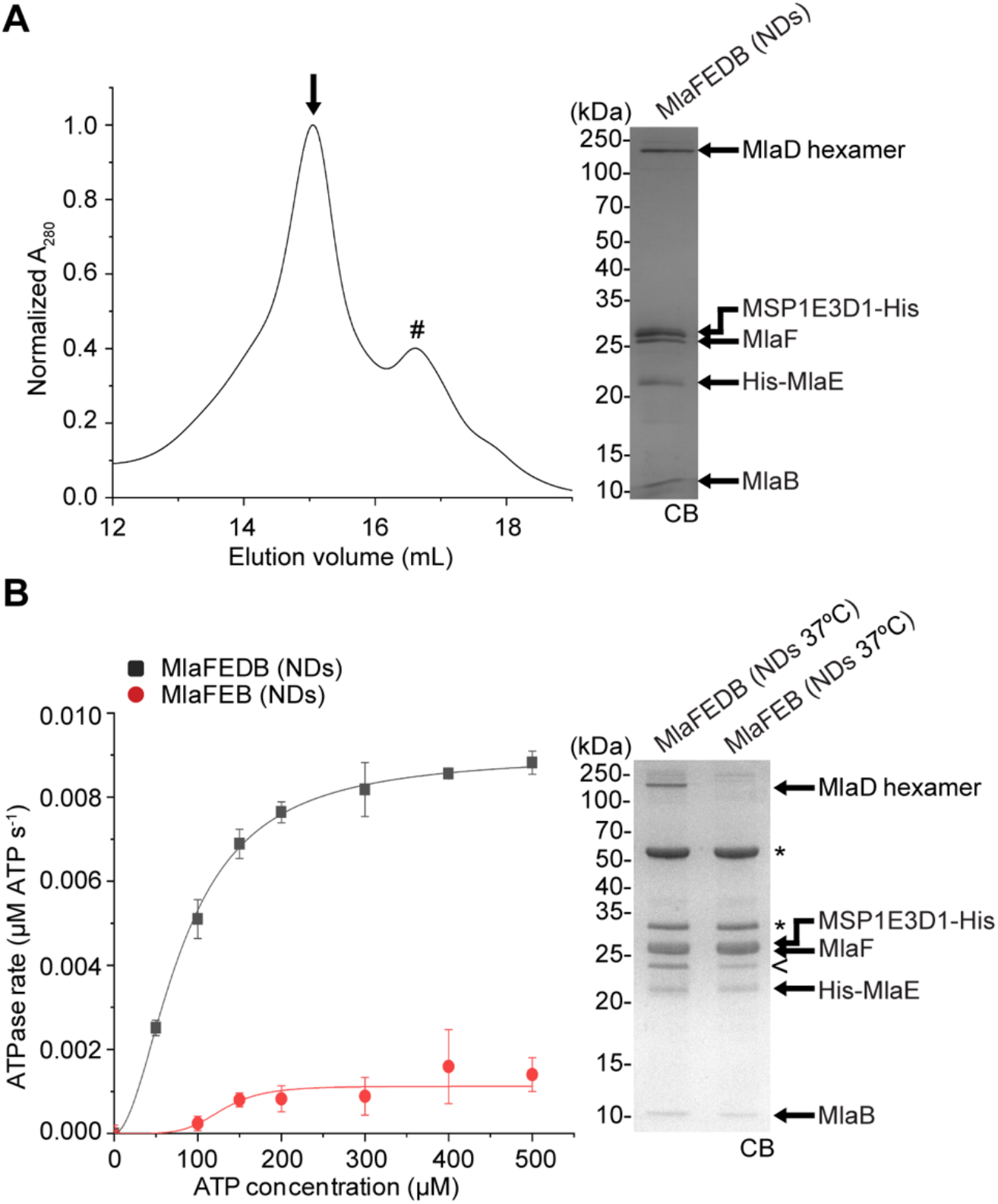
Nanodisc-embedded MlaFEDB exhibits higher ATP hydrolytic activity than MlaFEB. (A) Size-exclusion chromatographic profile of nanodisc-reconstituted (NDs) MlaFEDB complex. SDS-PAGE analysis of the peak fraction (annotated with an arrow (↓)) is shown on the right. #, empty NDs. (B) Enzyme-coupled ATPase assays of nanodisc-embedded MlaFEDB and MlaFEB complexes (0.1 μM) at 37°C. Average ATP hydrolysis rates from triplicate experiments were plotted against ATP concentrations, and fitted to an expanded Michaelis-Menten equation that includes a term for Hill coefficient (n); MlaFEDB NDs (k_cat_ = Vmax/[complex] = 0.090 ± 0.002 μmol ATP s^-1^/μmol complex, K_m_ = 82.5 ± 4.3 μM, n = 1.9 ± 0.2) and MlaFEB NDs (k_cat_ = 0.011 ± 0.002 μmol ATP s^-1^/μmol complex, K_m_ = 129.1 ± 12.8 μM, n = 5.3 ± 2.0). Errors depicted by bars and ± signs are standard deviations of triplicate data. SDS-PAGE analysis of the complexes used for these assays is shown on the right. *, pyruvate kinase/lactate dehydrogenase enzymes used in coupled assay; <, degraded MSP1E3D1. CB, Coomassie blue staining.

### MlaC transfers PLs to MlaFEDB in spontaneous and ATP-enhanced manners in vitro

We next tested the possibility of retrograde transport and investigated whether nanodisc-embedded MlaFEDB is able to extract and receive PLs from MlaC at 37°C. We incubated tag-less MlaC bound to [^14^C]-labelled PLs (Figure S2A and S2B) with His-tagged MlaFEDB complex in non-labelled nanodiscs in a 5:1 ratio, and monitored transfer of radioactive lipids following optimal nickel affinity-based separation of the two components (Figure 2A and S2C). The nanodisc-embedded MlaFEB complex lacking MlaD was used as control; with or without ATP, a small amount of radioactive PLs on MlaC was transferred to the MlaFEB nanodiscs and treated as ‘non-specific’ background (∼11%) (Figure 2B). Interestingly, we observed a substantial increase in spontaneous transfer of radioactive PLs from MlaC to nanodiscs when MlaD was present in the complex (∼28% with background subtracted) (Figure 2B). Following activation with ATP, even more PLs were transferred to the MlaFEDB complex (∼41% with background subtracted; ∼1.5 fold above the level without ATP) (Figure 2C). Importantly, both the spontaneous and ATP-enhanced gains in radioactive PLs on the nanodiscs were accompanied by corresponding losses from MlaC (albeit with ∼10-15% radioactivity unrecovered), overall indicating point-to-point PL transfer (Figure 2B). Of note, we observed reduced levels of PL transfer to MlaFEDB with more MlaC added (c.f. MlaC:MlaFEDB = 1:1 vs 5:1, Figure S3), possibly trending towards saturation. We have therefore reconstituted specific MlaD- and ATP-dependent retrograde PL transport from MlaC to the IM ABC transporter in vitro.

**Figure 2.**
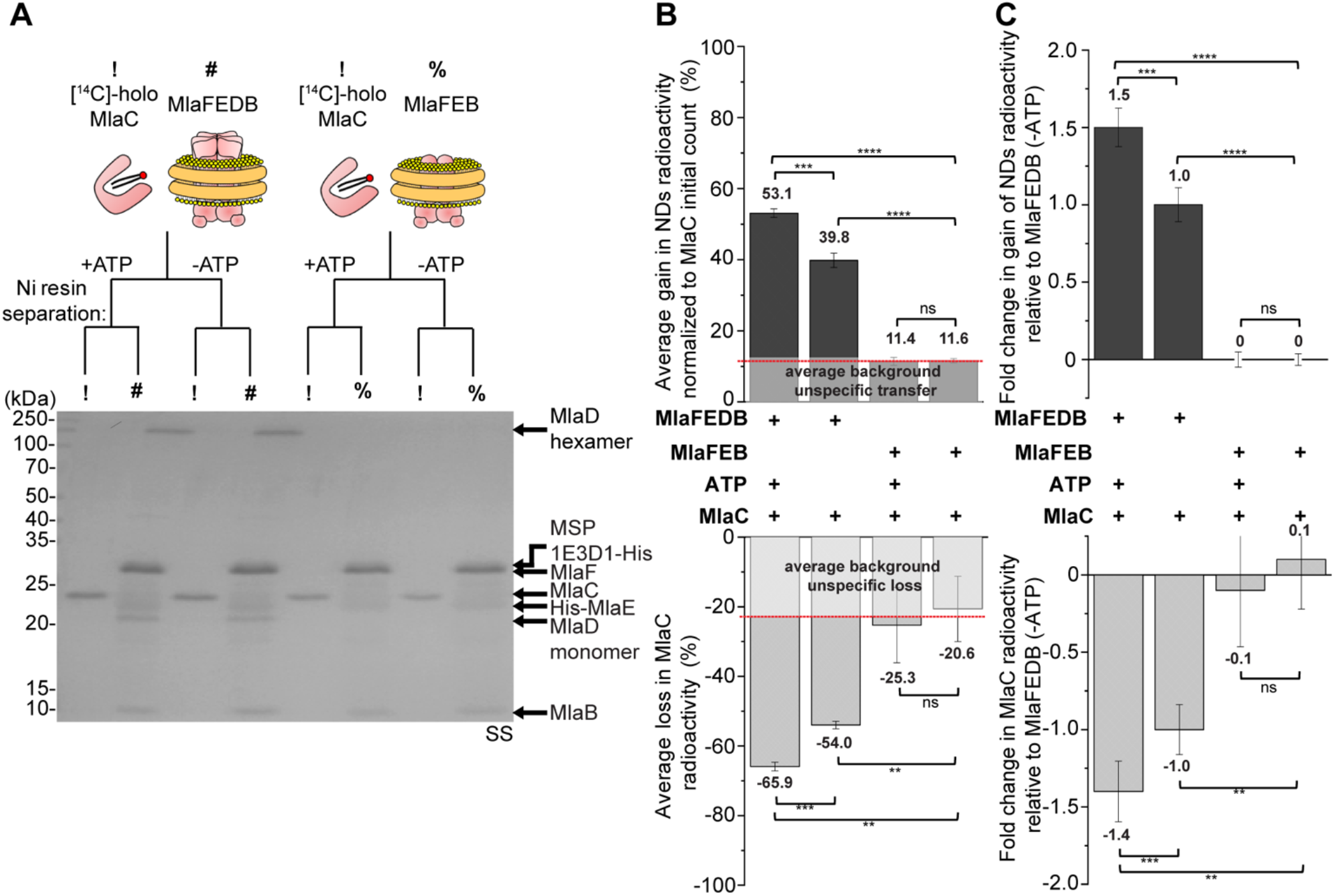
MlaC transfers [^14^C]-labelled lipids to nanodisc-embedded MlaFEDB in an MlaD- and ATP-dependent manner. (A) SDS-PAGE analysis of indicated samples following co-incubation with or without ATP, and subsequent nickel affinity-based clean separation of tag-less [^14^C]-PL bound (holo) MlaC (10 μM) and His-tagged non-radioactive nanodisc-embedded complexes (2 μM). !, [^14^C]-holo MlaC; #, nanodisc-embedded MlaFEDB; %, nanodisc-embedded MlaFEB; SS, silver staining. (B) Average gains and losses of radioactivity ([^14^C]-lipids) in the indicated nanodiscs (*top*) and MlaC (*bottom*), respectively, normalized to the initial counts on MlaC. Data from co-incubation of MlaFEB/MlaC are treated as unspecific background transfer/loss. (C) Fold changes in gains and losses of radioactivity in the indicated nanodiscs (*top*) and MlaC (*bottom*), respectively, derived from (B) after accounting for MlaFEB background and normalized to data from co-incubation of MlaFEDB/MlaC without ATP (MlaFEDB (-ATP)). Error bars represent standard deviations calculated from triplicate experiments. Student’s t-tests: ns, not significant; **, p < 0.01; ***, p < 0.001, ****; p < 0.0001.

### ATP hydrolysis by the MlaFEDB complex is coupled to retrograde PL transport

We next characterized the observed ATP-enhanced PL transfer from MlaC to nanodisc-embedded MlaFEDB. Mutations in K47 in the Walker A motif of MlaF and in the conserved T52 residue in MlaB have been reported to abolish ATP hydrolytic activity of the ABC transporter in detergents (18). We showed that the Walker A MlaF_K47A_ variant, which is unable to hydrolyze ATP in nanodiscs (Figure 3A and S4A), still allowed spontaneous PL transfer from MlaC to MlaF_K47A_EDB, but completely abolished ATP-enhanced transfer (Figure 3B and S4B). In addition, either the use of the non-hydrolyzable ATP analog, AMP-PNP (Figure 3C and S4C), or the ATPase inhibitor, vanadate (Figure 3D, S4D and S5), similarly led to the loss of ATP-activated PL transfer from MlaC to wildtype MlaFEDB, establishing that ATP hydrolysis is required for this process. Consistent with this idea, complexes with the MlaB_T52A_ variant (18), which we now show possesses partial ATPase activity in nanodiscs (Figures 3A and S4A), exhibited correspondingly lower ATP-enhanced PL transfer from MlaC (Figure 3B and S4B). We further demonstrated that the amount of radioactive PLs recovered on wildtype MlaFEDB nanodiscs reached a plateau with increasing concentrations of ATP (Figure 3E and S4E), establishing that ATP-activated PL transfer is saturable and thus specific. In fact, the K_M_ for ATP in enhanced PL transfer activity (33.1 ± 22.2 μM) is comparable to that in ATPase activity of the complex (70-83 μM across multiple experiments). We conclude that ATP hydrolysis is strongly coupled to retrograde PL transport mediated by the IM Mla complex.

**Figure 3.**
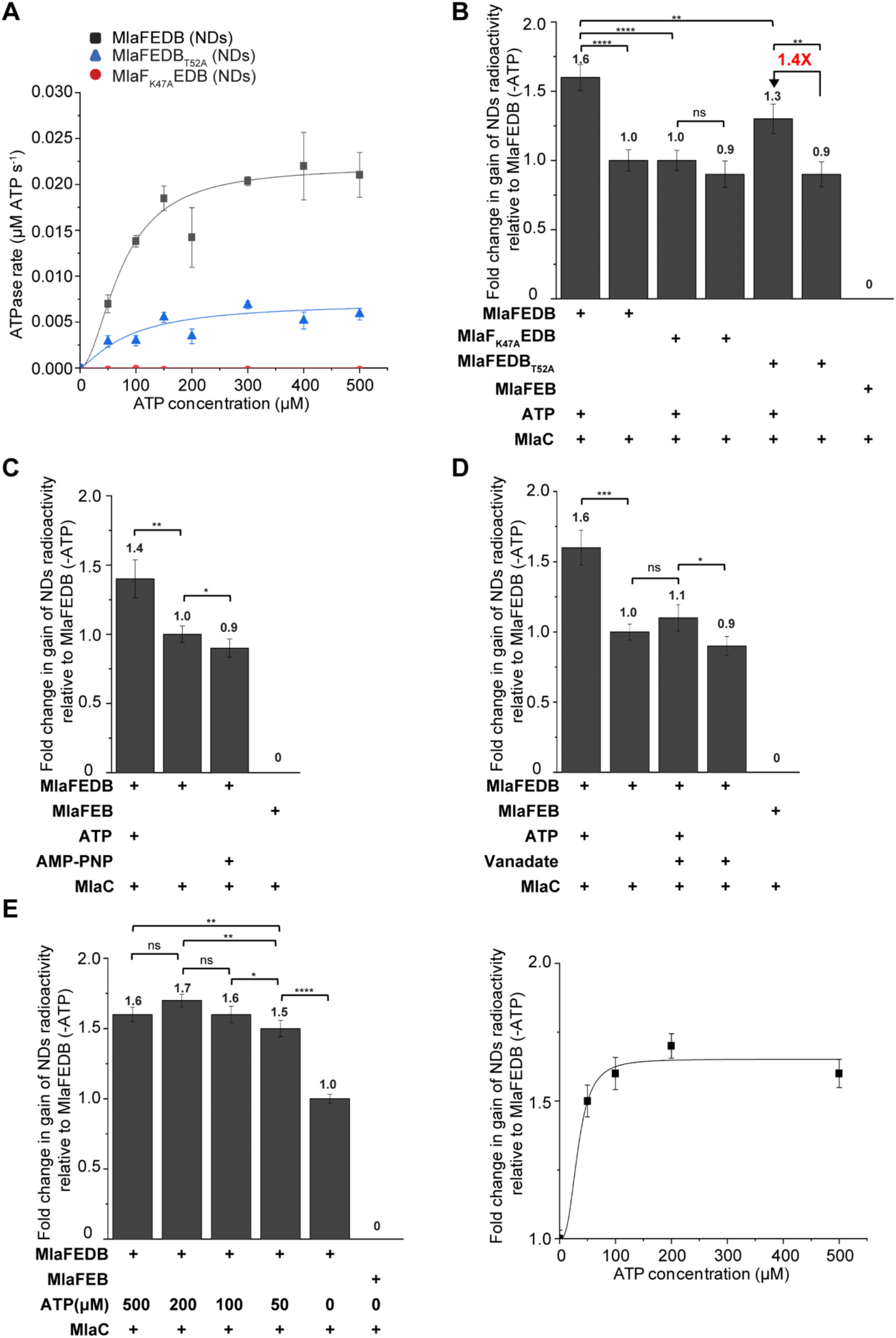
ATP hydrolysis is coupled to retrograde PL transport. (A) Enzyme-coupled ATPase assays of nanodisc-embedded MlaFEDB, MlaFEDB_T52A_ and MlaF_K47A_EDB complexes (0.1 μM) at 37°C. Average ATP hydrolysis rates from triplicate experiments were plotted against ATP concentrations, and fitted to an expanded Michaelis-Menten equation that includes a term for Hill coefficient (n); MlaFEDB NDs (k_cat_ = 0.220 ± 0.000 μmol ATP s^-1^/μmol complex, K_m_ = 75.0 ± 8.0 μM, n = 1.9 ± 0.4), MlaFEDB_T52A_ NDs (k_cat_ = 0.072 ± 0.028 μmol ATP s^-1^/μmol complex, K_m_ = 88.4 ± 62.1 μM, n = 1.3 ± 1.2). SDS-PAGE analysis of the complexes used for the assay is shown in Figure S4A. (B-E) Fold changes in gains of radioactivity ([^14^C]-lipids) in indicated nanodiscs, following co-incubation and subsequent nickel affinity-based separation of tag-less [^14^C]-PL bound (holo) MlaC (10 μM) and His-tagged non-radioactive nanodisc-embedded complexes (2 μM), for (B) ATPase mutant complexes, (C) addition of AMP-PNP, (D) addition of vanadate (2.7 mM), and (E) different ATP concentrations. Data from (E) are replotted on the right and fitted to an expanded Michaelis-Menten equation that includes a term for Hill coefficient (n); MlaFEDB NDs (K_m_ for ATP during PL transfer = 33.1 ± 22.2 μM, n = 2.8 ± 4.6, R^2^ = 0.9891). SDS-PAGE analyses depicting clean separation of nanodisc-embedded complexes and MlaC are shown in Figure S4B-E. Fold changes are derived from average gains in radioactivity counts in indicated nanodiscs, after accounting for MlaFEB background and normalized to data from MlaFEDB (-ATP). Errors depicted by bars and ± signs are standard deviations from triplicate data. Student’s t-tests: ns, not significant; *, p < 0.05; **, p < 0.01; ***, p < 0.001; ****, p < 0.0001.

### MlaD modulates PL binding capacity and ATP hydrolytic activity of the MlaFEDB complex in retrograde PL transfer

We next went on to investigate the role of MlaD, especially given that it is required for the transfer of PLs from MlaC to the MlaFEDB complex in the absence of ATP. To do that, we examined two non-functional MlaD variants that have previously been reported (MlaD_L106N/L107N_ and MlaD_Δ141-183_) (14). The L106N/L107N mutation is found at one of the constriction points of the central hydrophobic pore of MlaD hexamers, while the Δ141-183 deletion removes the C-terminal helices that form part of the pore; both are expected to affect lipid binding of the complex. In addition, MlaD_Δ141-183_ forms hexamers in the complex that are no longer SDS-resistant, exhibiting lower stability than those of wildtype MlaD and MlaD_L106N/L107N_ (Figure S6). Indeed, we observed elevated or reduced spontaneous transfer of radioactive PLs from MlaC to nanodisc-embedded MlaFED_L106N/L107N_B (∼1.3-fold) or MlaFED_Δ141-183_B (∼0.7-fold, compared to wildtype MlaFEDB), respectively (Figure 4A and S7A). As expected, MlaD directly modulates the PL binding capacity of the MlaFEDB complex.

**Figure 4.**
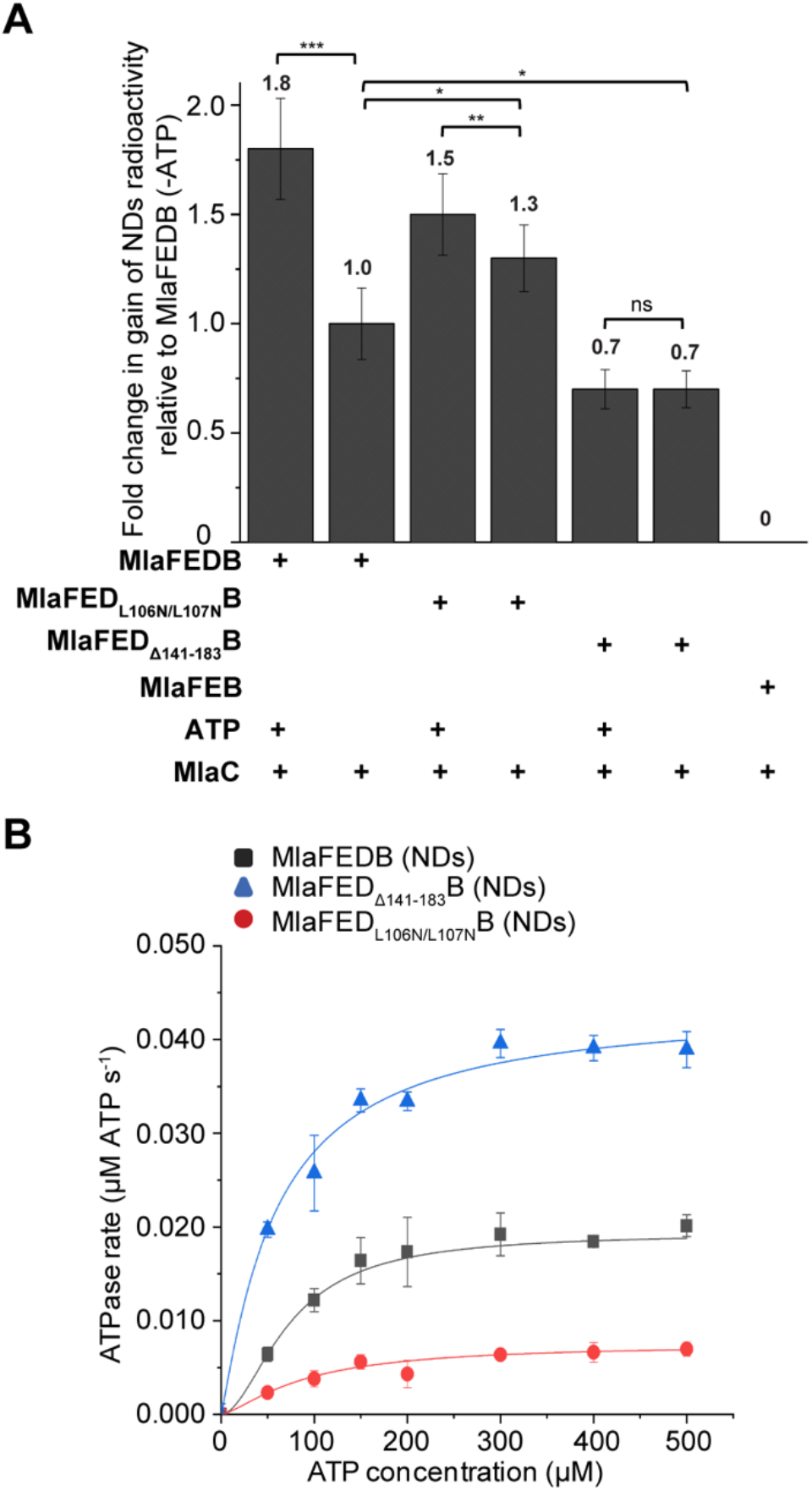
MlaD modulates both the spontaneous and ATP-activated PL transfer steps. (A) Fold changes in gains of radioactivity ([^14^C]-lipids) in indicated nanodiscs, following co-incubation and subsequent nickel affinity-based separation of tag-less [^14^C]-PL bound (holo) MlaC (10 μM) and His-tagged non-radioactive nanodisc-embedded complexes containing MlaD variants (2 μM). SDS-PAGE analyses depicting clean separation of nanodisc-embedded complexes and MlaC are shown in Figure S7A. Fold changes are derived from average gains in radioactivity counts in indicated nanodiscs, after accounting for MlaFEB background and normalized to data from MlaFEDB (-ATP). (B) Enzyme-coupled ATPase assays of nanodisc-embedded MlaFEDB, MlaFED_L106N/L107N_B and MlaFED_Δ141-183_B complexes (0.1 μM) at 37°C. Average ATP hydrolysis rates from triplicate experiments were plotted against ATP concentrations, and fitted to an expanded Michaelis-Menten equation that includes a term for Hill coefficient (n); MlaFEDB NDs (k_cat_ = 0.193 ± 0.007 μmol ATP s^-1^/μmol complex, K_m_ = 73.2 ± 6.9 μM, n = 1.8 ± 0.4), MlaFED_L106N/L107N_B NDs (k_cat_ = 0.075 ± 0.011 μmol ATP s^-1^/μmol complex, K_m_ = 88.7 ± 26.3 μM, n = 1.4 ± 0.5) and MlaFED_Δ141-183_B NDs (k_cat_ = 0.436 ± 0.027 μmol ATP s^-1^/μmol complex, K_m_ = 59.2 ± 7.5 μM, n = 1.1 ± 0.2). SDS-PAGE analysis of the complexes used for the assay is shown in Figure S7B. Errors depicted by bars and ± signs are standard deviations from triplicate data. Student’s t-tests: ns, not significant; *, p < 0.05; **, p < 0.01; ***, p < 0.001.

Interestingly, these MlaD mutations also affected the ATP hydrolytic activities of the corresponding complexes. The MlaFED_L106N/L107N_B complex exhibited reduced ATPase activity (Figure 4B and S7B) and hence had much lower ATP-enhanced PL transfer from MlaC (Figure 4A and S7A). Remarkably, the MlaFED_Δ141-183_B variant displayed much greater ATPase activity than wildtype complex (Figure 4B and S7B), yet was unable to activate additional PL transfer in the presence of ATP (Figure 4A and S7A). We conclude that MlaD modulates ATPase function and is essential for coupling ATP hydrolysis to PL transfer in the MlaFEDB complex.

### ATP hydrolysis prevents spontaneous transfer of PLs from MlaFEDB to MlaC

We have now established that MlaC spontaneously transfers PLs to the MlaFEDB complex in a manner that can be enhanced by ATP hydrolysis. However, passive transfer of PLs from MlaFEDB to MlaC has also been reported (31). To address this discrepancy, we also studied PL transfer in the anterograde direction using our nanodisc-reconstituted complexes. We reconstituted MlaFEDB and MlaFEB complexes into nanodiscs containing [^14^C]-labelled PLs (Figure S8), and incubated these purified nanodiscs with tag-less apo MlaC. As negative control, we showed that nanodiscs embedded MlaFEB transferred no or little PLs to MlaC (presumed non-specific transfer up to ∼5%) (Figure 5A and 5B). In the presence of MlaD, the MlaFEDB complex spontaneously transferred ∼20% of radioactive PLs from the nanodiscs to MlaC, supporting the possibility of anterograde transport (Figure 5A and 5C). Remarkably, however, addition of ATP entirely prevented transfer of PLs from MlaFEDB to MlaC, and this phenomenon was inhibited by addition of vanadate. Therefore, even though MlaD can spontaneously transfer PLs to MlaC, ATP hydrolysis by the IM complex abolishes such activity.

**Figure 5.**
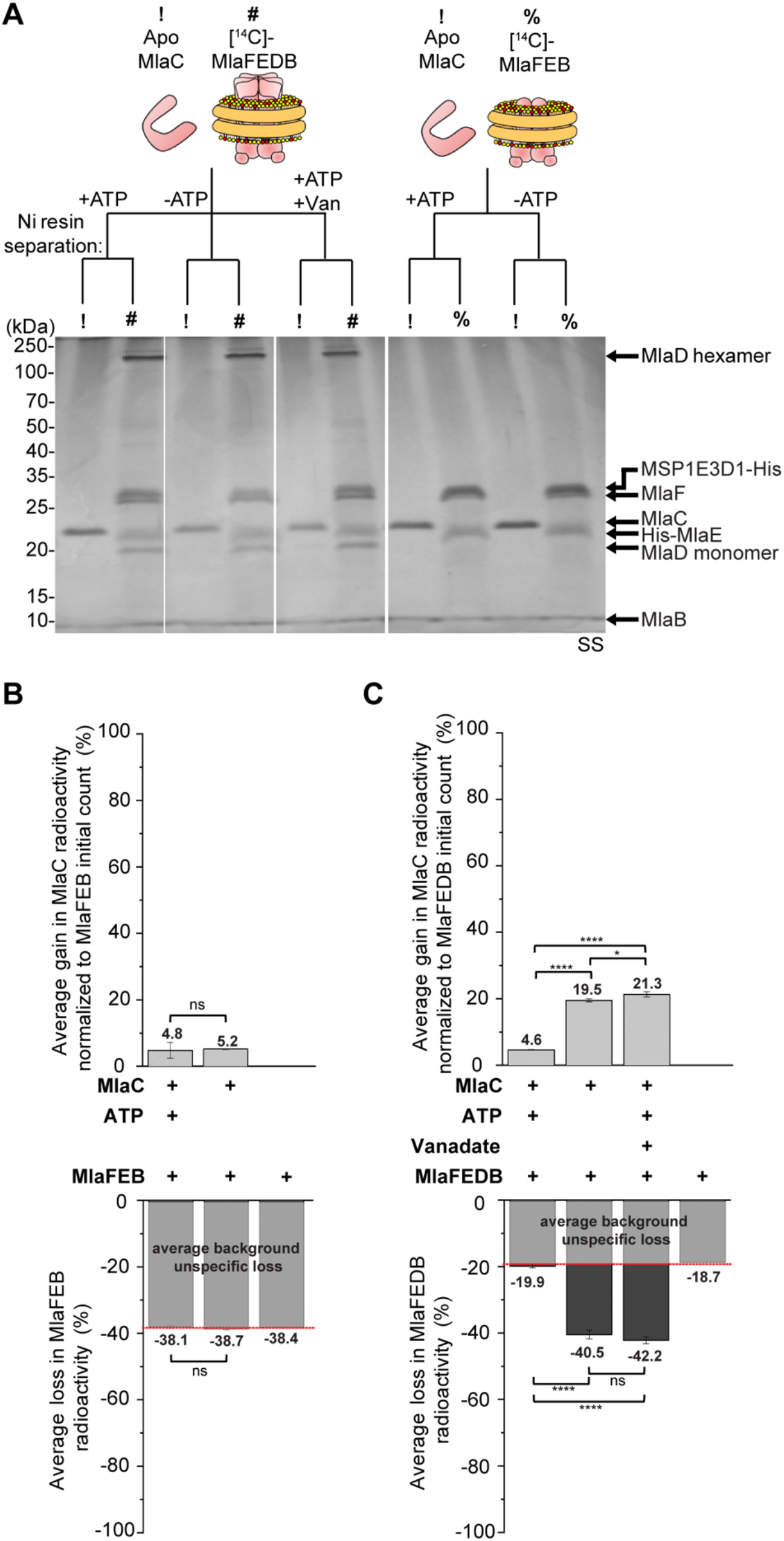
ATP hydrolysis abolishes spontaneous transfer of PLs from MlaFEDB to MlaC. (A) SDS-PAGE analysis of indicated samples following co-incubation with or without ATP, or vanadate, and subsequent nickel affinity-based clean separation of tag-less non-radioactive apo MlaC (10 μM) and His-tagged nanodisc-embedded complexes reconstituted with [^14^C]-PLs (2 μM). !, apo MlaC; #, nanodisc-embedded [^14^C]-MlaFEDB; %, nanodisc-embedded [^14^C]-MlaFEB; SS, silver staining. (B) Average gains and losses of radioactivity ([^14^C]-lipids) in MlaC (*top*) and MlaFEB (*bottom*), respectively, normalized to the initial counts on MlaFEB. (C) Average gains and losses of radioactivity ([^14^C]-lipids) in MlaC (*top*) and MlaFEDB (*bottom*), respectively, normalized to the initial counts on MlaFEDB. Data from co-incubation of MlaFEB/MlaFEDB with nickel resin alone are treated as unspecific background loss. Error bars represent standard deviations calculated from triplicate experiments. Student’s t-tests: ns, not significant; *, p < 0.05; ****; p < 0.0001.

## Discussion

In this work, we have reconstituted ATP-powered retrograde PL transport mediated by the IM ABC transporter of the OmpC-Mla system, using purified components. By applying [^14^C]-labelled lipids, we have demonstrated spontaneous PL transfer from MlaC to the MlaFEDB complex, with enhanced lipid transfer occurring upon ATP hydrolysis (Figure 2). We have also observed spontaneous PL transfer from MlaFEDB to MlaC but this transfer is abolished in the presence of ATP (Figure 5). Therefore, we have shown that PLs equilibrate between MlaC and MlaFEDB, and ATP hydrolysis drives overall transport in the retrograde direction. Interestingly, the fold of the MlaE permease domains resemble that of the LptFG counterparts in the LPS exporter (19), yet the MlaFEDB complex possesses lipid import function; such ABC importers with ‘exporter’ folds have only recently been discovered (33-35).

Here is a mechanistic model for MlaFEDB function, integrating all available biochemical and structural information (Figure 6). In the first step, PL-bound MlaC arrives at the IM, and interacts with the MlaFEDB complex via the protomer interfaces of MlaD hexamers (15). This ensues the reversible partitioning of the PL molecule(s) between the lipid binding pockets of MlaC and the complex, even in the absence of ATP. While MlaD hexamers alone bind PLs much less strongly than MlaC (15), the MlaFEDB complex, where several structures now reveal a combined lipid binding cavity formed by both MlaD and MlaE (19-23), must possess strong affinity for PLs, likely comparable to MlaC. The trajectory of PL movement between the two binding sites remains to be clarified. In the second step, ATP binding has been shown to induce two conformational states of the MlaFEDB complex, believed to be sequential in the process (21). Initial ATP binding to MlaF appears to cause a conformational change involving upward movement of MlaD away from MlaE, which likely releases the PL molecule(s) from MlaD binding to be eventually bound by MlaE alone. Subsequent complete binding of ATP induces tight dimerization of MlaF, in turn collapsing the MlaE lipid binding cavity (21, 23), eventually forcing bound PL molecule(s) into the membrane. Whether PLs enter the outer leaflet of the IM, or get flipped across the bilayer, is not known. In the final step, ATP hydrolysis resets the MlaFEDB complex, allowing it to again adopt the nucleotide-free state. ATP binding and/or hydrolysis are therefore required to overcome the energy barrier conferred by tight binding of PLs within the lipid binding cavity of the MlaFEDB complex, and force them into the IM. Supporting the idea that ATP is not instead needed to release lipids directly from MlaC, MlaC itself is unable to activate ATPase activity of the complex (Figure S1B). This whole ATP-driven process ultimately favors retrograde PL transfer from MlaC, via the MlaFEDB complex, to the IM, in accordance with the Le Chatelier’s principle.

**Figure 6.**
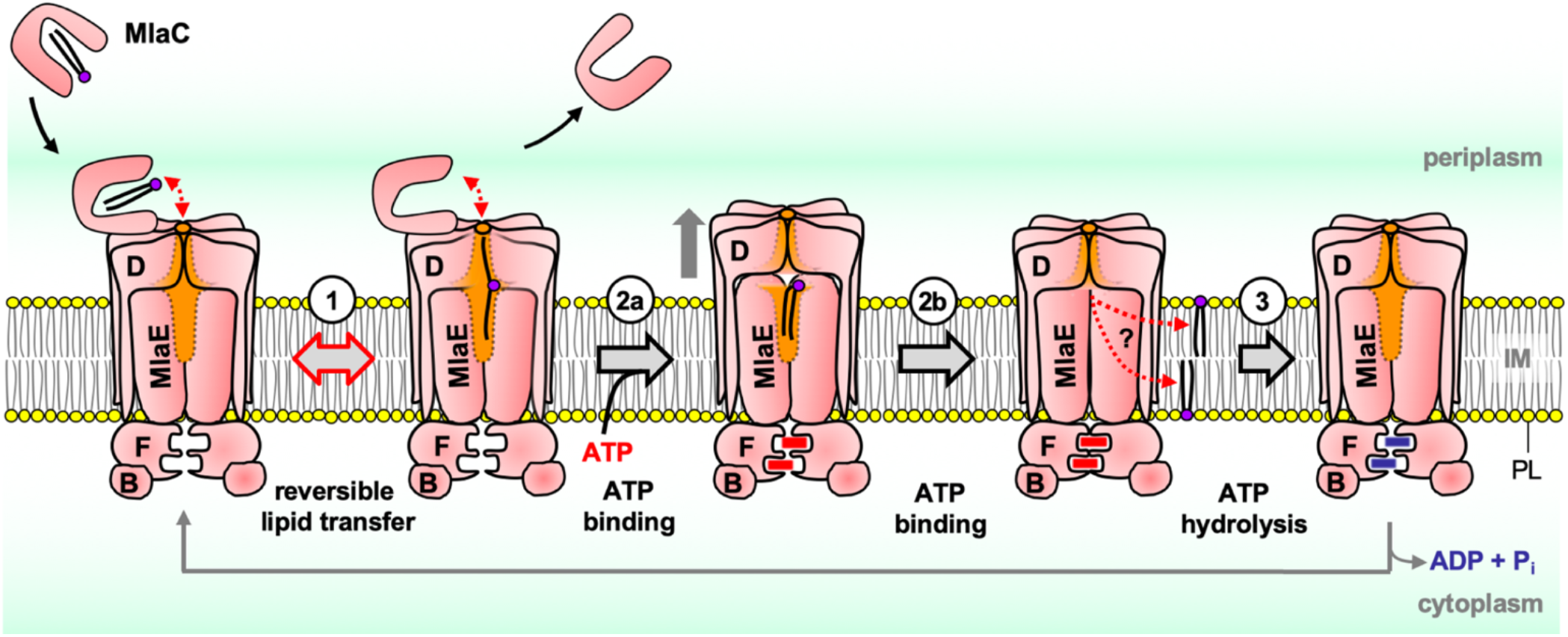
Proposed mechanistic model for MlaFEDB function in retrograde PL transfer. Step 1: Upon MlaC binding, reversible spontaneous transfer of PL cargo occurs between the lipid binding pocket in MlaC and the lipid binding cavity formed by both MlaD and MlaE in the complex (illustrated in orange). Step 2a: Initial ATP binding to MlaF may induce a state where MlaD moves away from MlaE (illustrated by grey arrow), possibly facilitating lipid binding by MlaE alone (21). Step 2b: Complete ATP binding eventually causes dimerization of MlaF, along with the collapse of the lipid binding cavity in MlaE, thus forcing the PL molecule into the membrane (21, 23). It is not clear which leaflet of the IM the PL molecule goes into. Step 3: Hydrolysis of ATP then resets such conformational changes, so the complex can return to the nucleotide apo state.

In the context of the model, MlaD influences both the spontaneous and ATP-coupled steps of PL transfer. We speculate that the conformational flexibility of MlaD hexamers plays a major role in achieving such modulation. Unlike wildtype MlaD, the MlaD_Δ141-183_ variant exists as hexamers in the complex that are not stable to SDS (Figure S6), representing a “loose” form of the hexamer with greater conformational freedom. In support of this idea, the soluble domain of this variant can form heptamers in vitro (14). The MlaFEDB complex with a “loose” MlaD hexamer would have a more flexible lipid binding cavity, therefore incurring greater entropic costs to bind ligands. The MlaFED_Δ141-183_B complex may thus have a lower affinity for PLs, resulting in reduced spontaneous PL transfer from MlaC (Figure 4). Increased conformational flexibility may also account for the much higher ATPase activity of this variant complex, possibly by facilitating conformational movements in the MlaE permease domains associated with the ATP hydrolytic cycle; however, this led to uncoupling between ATP hydrolysis and PL transfer. By analogy, we posit that the MlaD_L106N/L107N_ variant forms “tight” hexamers in the complex, perhaps due to more hydrogen bonding interactions in the central pore. Decreased conformational flexibility thereby confers an increase in affinity for spontaneous PL binding and yet a reduced ATPase activity (Figure 4). Ultimately, the MlaFED_L106N/L107N_B complex has much lower ATP-enhanced PL transfer from MlaC, suggesting this non-functional variant is ineffective at forcing PLs out of the MlaDE cavity into the membrane, thus unable to drive overall flux of PLs from the OM to the IM. Overall, we propose that the conformational flexibility in the wildtype MlaD hexamer is optimally tuned to strike a balance between PL binding and ATPase activity, yet ensure productive coupling between PL transfer and ATP hydrolysis. Still, MlaD mutations may also have direct impact on PL binding enthalpies and/or interactions with MlaC. Further studies to test these ideas would be needed.

This reconstitution addresses the controversy of directionality, and establishes that ATP drives retrograde PL transport mediated by the OmpC-Mla system. Our work thus validates the interpretation of OmpC-Mla function in existing genetic studies (10, 11, 26, 27), and is consistent with plastid or mycobacterial homologs of the Mla IM complex being characterized as lipid uptake systems (36-38). Our highly sensitive PL transfer assay demonstrates point-to-point movement of native [^14^C]-labelled PLs between MlaC and MlaFEDB, and reveals both reversible spontaneous and irreversible ATP-activated steps in the process. In the midst of preparing this manuscript, Tang et al. have independently developed an indirect assay using non-native fluorescent lipids, and similarly reported that the Mla pathway mediates ATP-dependent retrograde PL transport in vitro (20). While our assays are necessarily complementary, the direct nature of the PL transfer assay developed here, and the use of native lipids, ultimately provide stronger inference about corresponding events happening in the bacterial cell. Importantly, our work has additionally uncovered the existence of a spontaneous retrograde PL transfer step from MlaC to MlaFEDB, which was not detected in the Tang et al. (20) or earlier studies (31). We have also observed spontaneous anterograde PL transfer from MlaFEDB to MlaC; contrary to these previous studies (20, 31), however, we have demonstrated that such anterograde transfer is not ATP-independent, but is abolished and prevented by ATP hydrolysis. These two significant findings set us apart from the Tang et al. work, directly advancing our understanding of how ATP binding/hydrolysis is coupled to PL transfer at the MlaFEDB complex to ultimately drive PL transport in the retrograde fashion. Collectively, both the fluorescence-based and radioactivity-based assays establish the directionality of the system. These PL transport assays will continue to serve as critical tools for detailed mechanistic studies, and for future identification and validation of small molecule inhibitors against the OmpC-Mla pathway. Adapted assays can also be applied to study possible lipid transport mediated by other putative systems, including the Tol-Pal complex (30) and other MCE domain-containing complexes (14, 39, 40).

## Materials and Methods

### Reconstitution of membrane proteins in lipid nanodiscs

The reconstitutions of purified membrane proteins (MlaF(His-E)B, MlaF(His-E)DB, MlaF_K47A_(His-E)DB, MlaF(His-E)DB_T52A_, MlaF(His-E)D_L106/L107N_B and MlaF(His-E)D _Δ141-183_B) into nanodiscs were adapted from published protocols (32). 10 mg of *E. coli* polar lipid extracts (Avanti Polar Lipids) were dissolved in 1 mL of chloroform and dried overnight. 1 mL of TBS buffer (20 mM Tris.HCl pH 8.0, 150 mM NaCl) containing 25 mM sodium cholate (Sigma-Aldrich) was added to the dried lipid film, and vortexed and sonicated until a clear solution was obtained. The respective Mla complex, the membrane scaffold protein MSP1E3D1-His and lipid were mixed at a molar ratio of 1:10:1500 in TBS and incubated, with rocking, for 1 h at 4 °C. Bio-beads SM2 resin (Bio-Rad) was subsequently added to the solution (30 mg per 1-mL reconstitution mixture) and incubated with gentle agitation for 3 h at 4 °C. Upon removal of Bio-beads, the sample was concentrated and purified by size exclusion chromatography (SEC) on a Superose 6 Increase 10/300 GL column (GE Healthcare) on ÄKTA™ Pure. 0.5-mL fractions were collected and subjected to SDS-PAGE analysis and Coomassie blue staining (InstantBlue™, expedeon). Fractions containing the nanodisc-embedded complex were pooled and concentrated on a 30 kDa cut-off ultra-filtration device (Amicon Ultra, Merck Millipore). For the reconstitution of radioactive nanodisc-embedded MlaF(His-E)B and MlaF(His-E)DB complexes, [^14^C]-labelled PLs extracted from [^14^C]-acetate labelled cells were used in place of *E. coli* polar lipid extracts, and the nanodiscs samples were purified on a self-packed column containing Superose 6 Prep Grade (GE Healthcare) instead.

### In vitro PL transfer assays

For in vitro retrograde PL transfer experiments, 10 μM of purified tag-less MlaC bound to [^14^C]-labelled PLs was incubated with 2 μM of purified non-radioactive His-tagged nanodisc-embedded complex. For in vitro anterograde PL transfer experiments, 10 μM of purified tag-less apo MlaC was incubated with 2 μM of purified His-tagged nanodisc-embedded complex reconstituted with [^14^C]-labelled PLs. These components were incubated in 400 μL TBS buffer, either in the absence or presence of 2 mM ATP (Sigma-Aldrich) and 2 mM MgCl_2_, for 10 min at 37°C. After the transfer reaction, the mixture was topped up with TBS to a final volume of 2.5 mL, and incubated with 0.5 mL nickel resin (His60 Ni Superflow Resin, Takara Bio) in a Poly-Prep chromatography column (BioRad) for 1h, with rocking, at room temperature. The filtrate (containing separated tag-less MlaC) was collected after the mixture was passed through the resin twice. The resin was washed with 5 × 4 mL TBS containing 30 mM imidazole, and the nanodisc-embedded complex was eluted with 2.5 mL TBS containing 300 mM imidazole. Samples obtained from the nickel affinity-based separation were subjected to SDS-PAGE analysis and visualized by silver staining to ensure that there was minimal cross-contamination of the two components. For experiments involving vanadate, 2 μM of the nanodisc-embedded MlaFEDB complex was first pre-incubated with 2 mM ATP and 2 mM MgCl_2_ for 5 min, then with 2.7 mM vanadate for 10 min, before finally incubating with 10 μM of MlaC for 10 min for PL transfer. 2 mM AMP-PNP (Sigma-Aldrich) was also used in place of ATP for specific transfer experiments. All transfer experiments were conducted in triplicates.

After separation of the filtrate (tag-less MlaC) and eluate (His-tagged nanodisc-embedded complex), 2 mL of each component was added to 2 mL Ultima Gold scintillation fluid (PerkinElmer), and radioactivity counted on the MicroBeta2 scintillation counter (PerkinElmer). The recovery of tag-less MlaC was quantified using the ratio standard curve method (41) while that of His-tagged nanodisc-embedded complexes were quantified using the Lowry assay (DC Protein Assay, Bio-Rad). For retrograde assay, the average gains and losses of radioactivity in the nanodiscs and MlaC, respectively, were normalized to the initial [14C]-counts in MlaC using the following equation:

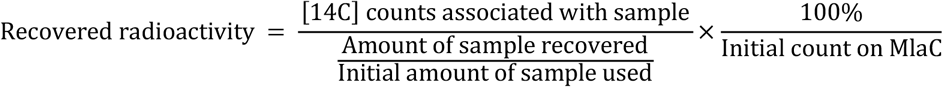

Fold changes in gains and losses of radioactivity in the nanodiscs and MlaC, respectively, were derived after accounting for MlaFEB background and normalized to data from co-incubation of MlaFEDB/MlaC without ATP (MlaFEDB (-ATP)). The starting radioactivity on [^14^C]-PL-bound MlaC were in the range of 12790-22854 cpm (or 13606-24313 dpm).

For the anterograde assay, the average gains and losses of radioactivity in the MlaC and nanodiscs, respectively, were normalized to the initial [14C]-counts in nanodisc using the following equation:

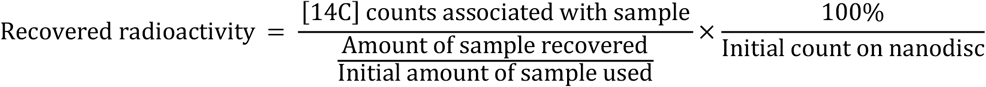

The starting radioactivity on nanodiscs containing [^14^C]-PLs were in the range of 75485-95725 cpm (or 80303-101835 dpm).

## Acknowledgments

W.-Y.L. was supported by the National University of Singapore President’s Graduate Fellowship (PGF). This work was supported by the Singapore Ministry of Health National Medical Research Council under its Open Fund Individual Research Grant (MOH-000145) (to S.-S.C.). The authors declare no conflict of interest.

