## Supplementary Information for "ATP disrupts lipid binding equilibrium to drive retrograde transport critical for bacterial outer membrane asymmetry"

**This PDF file includes:**

Supplementary Materials and Methods

Figures S1 to S8

SI References

### Supplementary Materials and Methods

#### Overexpression and purification of tag-less MlaC bound to [$^{14}$C]-labelled lipids

His-MlaC was over-expressed and purified from BL21($\lambda$DE3) cells harboring pETMHis-MlaC, which encodes MlaC without its signal sequence but with an N-terminal $His_6$ tag with a thrombin cleavage site (1). A 3-mL culture was grown from a single colony in LB broth supplemented with 200 µg/mL ampicillin (Sigma-Aldrich) and 1.25 µCi/mL [1-$^{14}$C]-acetate (PerkinElmer, NEC084A001MC) at 37 °C until $OD_{600}$ ~ 0.6. The cell culture was then used to inoculate into 150 mL LB broth containing 1.25 µCi/mL [$^{14}$C]-acetate and grown at the same temperature until $OD_{600}$ ~ 0.6. 1 mM IPTG (Axil Scientific, Singapore) was added and the culture was grown at 37 °C for another 3 h. Cells were pelleted by centrifugation at 4,700 x *g* for 20 min using swinging buckets and then resuspended in 8 mL buffer (20 mM Tris.HCl pH 8, 300 mM NaCl, 5 mM imidazole). Buffers were supplemented with 1 mM PMSF (Calbiochem), 50 µg/mL DNase I (Sigma-Aldrich) and 100 µg/mL lysozyme (Calbiochem). Cells were passed twice through a high-pressure French Press (French Press G-M, Glen Mills) homogenizer at 20,000 psi. Cell debris was removed by centrifugation at 4,700 x *g* for 10 min at 4 °C. Subsequently, supernatant was subjected to ultra-centrifugation (Model Optima L-100K, Beckman Coulter) on a SW 41 Ti rotor at 145,000 x *g* for 1 h at 4 °C to separate membrane and soluble fractions. Soluble fraction was incubated with 2.5 mL His60 Ni Superflow Resin (Takara Bio Inc) and rocked for 1 h on ice. The mixture was later loaded onto a column and allowed to drain by gravity. The filtrate was passed through the resin again, drained and the column was washed with 5 x 20 mL of wash buffer (20 mM Tris.HCl pH 8.0, 300 mM NaCl, 20 mM imidazole) and eluted with 8 mL of elution buffer (20 mM Tris.HCl pH 8.0, 150 mM NaCl, 200 mM imidazole). The eluate was subjected to at least five rounds of buffer exchange centrifugation at 4,000 x *g* with 10 mL TBS (20 mM Tris.HCl pH 8.0, 150 mM NaCl) by using an Amicon Ultra 10 kDa cut-off ultra-filtration device (Merck Millipore). The solution was then concentrated in a new 10 kDa cut-off ultra-filtration device (by centrifugation at 4,000 x *g* to ~500 µL. Tag-less MlaC was prepared by cleaving the N-terminal His-tag with the addition of thrombin (1 U/0.3 mg His-MlaC) and incubated at 22 °C overnight. After overnight digestion, protease activity was stopped by adding 2 mM protease inhibitor, PMSF. The tag-less MlaC was eventually isolated as the filtrate following incubation and rocking for 1 h on ice with 2.5 mL His60 Ni Superflow Resin (Takara Bio Inc).

#### Radioactive lipid extraction and thin layer chromatography

[$^{14}$C]-labelled PLs were obtained by extracting lipids from membrane fractions that have been metabolically labelled with [$^{14}$C]-acetate similar to how His-MlaC was over-expressed. 1 mg/100 µL of [$^{14}$C]-labelled holo MlaC protein or 1 mL of radiolabeled membrane fraction were used for the PL extraction according to the Bligh-Dyer method (2). Purified MlaC or the radiolabeled membrane fraction were mixed with 3.75 volumes of chloroform:methanol:TBS (1:2:0.8 vol/vol). The mixtures were vortexed and sonicated sequentially for 30 seconds for three times. The mixtures were centrifuged at 21,000 x *g* for 5 min, and the supernatant was recovered. 1.25 volumes of chloroform and 1.25 volumes of TBS were added to the supernatants. The mixtures were then centrifuged at 4000 x *g* for 5 min to separate organic and aqueous phases. The organic phase was gently removed to another vial, and the organic solvent was evaporated under $N_2$ gas. The dried lipids were dissolved in 10 uL chloroform:methanol (4:1 vol/vol) and loaded onto a TLC Silica gel 60 $F_{254}$ plate (Merck), if necessary. The plate was developed by the chloroform:methanol:water (65:25:4) solvent system, left to dry at room temperature, and visualized by phosphor imaging (STORM, GE healthcare).

**Overexpression and purification of tag-less apo MlaC**

The purification of apo MlaC was adapted from published protocols (1). 750 mL cell culture pellet from the growth of BL21(λDE3) cells harboring pETMHis-MlaC, was resuspended in 20 mL of lysis buffer (20 mM Tris.HCl pH 8, 300 mM NaCl and 8 M urea). Resuspended cells were rocked for 1.5 h at room temperature. The cells were lysed with three rounds of sonication on ice (30% power, 5-s pulse on, 5-s pulse off for 3 min). Cell debris was removed by centrifugation at 4,700 x *g* for 10 min at 4 °C. Subsequently, the supernatant was subjected to ultra-centrifugation (Model Optima L-100K, Beckman Coulter) at 145,000 x *g* for 1 h at 4 °C to separate membrane and soluble fractions. The supernatant was incubated with 2.5 mL His60 Ni Superflow Resin (Takara Bio Inc) and rocked for 2 h at room temperature. The resin mixture was later loaded onto a column and allowed to drain by gravity. The filtrate was passed through the resin again, drained and the column was washed with 2 x 15 mL of wash buffer 1 (20 mM Tris.HCl pH 8, 300 mM NaCl, 8 M urea and 1% SDS), 2 x 15 mL of wash buffer 2 (20 mM Tris.HCl pH 8, 300 mM NaCl, 8 M urea) and eluted with 10 mL of elution buffer (20 mM Tris.HCl pH 8.0, 150 mM NaCl, 8 M urea and 500 mM imidazole). The eluate was pipetted into Spectra/Por 3, 3.5 kDa MWCO, 18 mm flat width dialysis membranes and dialyzed against TBS for 2 h and then overnight. Next morning, the solution was transferred to a 10 kDa cut-off ultra-filtration device (Amicon Ultra, Merck Millipore) and concentrated by centrifugation at 4,000 x *g* to ~500 µL. Tag-less MlaC was prepared by cleaving the N-terminal His-tag with the addition of thrombin (1 U/0.3 mg His-MlaC) and incubated at 22 °C overnight. After overnight digestion, protease activity was stopped by adding 2 mM protease inhibitor, PMSF. The tag-less MlaC was eventually isolated as the filtrate following incubation and rocking for 1 h on ice with 2.5 mL

His60 Ni Superflow Resin (Takara Bio Inc). The filtrate was transferred to a 10 kDa cut-off ultra-filtration device (Amicon Ultra, Merck Millipore) and concentrated by centrifugation at 4,000 x *g* to ~500 µL. The protein was further purified by SEC system (AKTA, GE Healthcare) at 4 °C on a prepacked Superdex 200 increase 10/300 GL column using TBS, as the running buffer.

**Overexpression and purification of membrane protein complexes**

The purification of MlaFEDB/MlaFEB complexes was adapted from published protocols (3). MlaF(His-E)DB, $MlaF_{K47A}$(His-E)DB, MlaF(His-E)$D_{L106N/L107N}$B and MlaF(His-E)$D_{\Delta 141-183}$B were over-expressed and purified from BL21(λDE3) cells harboring pET22/42*mlaF(His-E)DCB*, pET22/42*mlaF*$_{K47A}$*(His-E)DCB*, pET22/42*mlaF(His-E)D*$_{L106N/L107N}$*CB* and pET22/42*mlaF(His-E)D*$_{\Delta 141-183}$*CB*, respectively. In order to optimize amounts of MlaB during the complex purification, a second over-expression vector pCDF*mlaB* was introduced into BL21((λDE3) cells. To overexpress MlaF(His-E)$DB_{T52A}$, pCDF*mlaB*$_{T52A}$ together with pET22/42*mlaF(His-E)DCB*$_{T52A}$ were introduced into BL21(λDE3) cells. To over-express MlaF(His-E)B, pCDF*mlaB* together with pET22/42*mlaF(His-E)* were introduced into BL21(λDE3) cells. A 30-mL culture was grown from a single colony in LB broth supplemented with 200 µg/mL ampicillin (Sigma-Aldrich) and 50 µg/ mL streptomycin (Sigma-Aldrich) at 37 °C until $OD_{600}$ ~ 0.6. The cell culture was then used to inoculate a 3-L culture and grown at the same temperature until $OD_{600}$ ~ 0.6. For MlaF(His-E)DB and associated variant complexes, 1 mM IPTG (Axil Scientific, Singapore) was added and the culture was grown at 37 °C for another 3 h. For MlaF(His-E)B complex, 0.1 mM IPTG was added and the culture was grown at 18 °C for another 20 h. Cells were pelleted by centrifugation at 4700 x g for 20 min and then resuspended in 25 mL buffer (20 mM Tris.HCl pH 8, 300 mM NaCl) buffer containing 1 mM PMSF (Calbiochem), 50 mg/mL DNase I (Sigma-Aldrich) and 100 mg/mL lysozyme (Calbiochem). Cells were passed twice through a high pressure French Press (French Press G-M, Glen Mills) homogenizer at 20,000 psi. Cell debris was removed by centrifugation at 4700 x *g* for 10 min at 4 °C. Subsequently, the supernatant was subjected to ultra-centrifugation (Model Optima L-100K, Beckman Coulter) at 145,000 x *g* for 1 h at 4 °C to separate membrane and soluble fractions. The membrane pellet fraction was extracted (20 mL of 20 mM Tris.HCl pH 8.0, 300 mM NaCl, 5 mM $MgCl_2$, 10% glycerol, 1% n-dodecyl b-D-maltoside (DDM) (Merck Millipore), 10 mM imidazole and 1 mM PMSF) and subjected to a second round of ultra-centrifugation at 145,000 x *g* for 1 h at 4 °C. The supernatant was incubated with 2.5 mL His60 Ni Superflow Resin (Takara Bio Inc) and rocked for 2 h on ice. The mixture was later loaded onto a column and allowed to drain by gravity. The filtrate was passed through the resin again, drained and the column was washed with 10 x 10 mL of wash buffer (20 mM Tris.HCl pH 8.0, 300 mM NaCl, 20 mM imidazole, 1mM PMSF, 5 mM $MgCl_2$, 10% glycerol and 0.05% DDM) and eluted with 15 mL of elution buffer (20 mM Tris.HCl pH 8.0, 150 mM NaCl, 200 mM imidazole, 1mM PMSF, 5 mM $MgCl_2$, 10% glycerol and 0.05% DDM). The eluate was concentrated in an Amicon Ultra 10 kDa cut-off ultra-filtration device (Merck Millipore) by centrifugation at 4,000 x *g* to ~500 µL. Proteins were further purified by size-exclusion chromatography (SEC) system (AKTA, GE Healthcare, UK) at 4 °C on a pre-packed Superdex 200 increase 10/300 GL column, using 20 mM Tris.HCl pH 8.0, 150 mM NaCl, 5 mM $MgCl_2$, 10% glycerol and 0.05% DDM as the eluent. For MlaF(His-E)B complexes, two columns were connected in series to allow better peak separation.

**Overexpression and purification of membrane scaffold protein MSP1E3D1-His**

The purification of MSP1E3D1-His was adapted from published protocols (4). MSP1E3D1 nanodiscs (~12.9 nm in diameter) were chosen because measured ATPase activities of MlaFEDB complexes (~11 nm in diameter) reconstituted in MSP1E3D1 or larger MSP2N2 (15.0-16.5 nm in diameter) nanodiscs did not exhibit any distinct differences, indicating that the size of the MSP1E3D1 nanodisc is sufficient in the context of ATP hydrolysis and PL transfer. MSP1E3D1-His was over-expressed and purified from BL21(λDE3) cells harboring pMSP1E3D1. A 15-mL culture was grown from a single colony in LB broth supplemented with 25 µg/mL kanamycin (Sigma-Aldrich) at 37 °C until $OD_{600}$ ~ 0.6. The cell culture was then used to inoculate a 1.5-L culture and grown at the same temperature until $OD_{600}$ ~ 0.6. 1 mM IPTG (Axil Scientific, Singapore) was added and the culture was grown at 37 °C for another 3 h. Cells were pelleted by centrifugation at 4700 x *g* for 20 min and then resuspended in 20 mL of 20 mM phosphate buffer pH 7.4 buffer containing 1 mM PMSF (Calbiochem), 50 mg/mL DNase I (Sigma-Aldrich) and 1% Triton X-100 (Sigma-Aldrich). Cells were passed twice through a high pressure French Press (French Press G-M, Glen Mills) homogenizer at 20,000 psi. Cell debris was removed by centrifugation at 4700 x *g* for 10 min at 4 °C. Subsequently, the supernatant was subjected to ultra-centrifugation (Model Optima L-100K, Beckman Coulter) at 145,000 x *g* for 1 h at 4 °C to separate membrane and soluble fractions. Soluble fraction was incubated with 2.5 mL His60 Ni Superflow Resin (Takara Bio Inc) and rocked for 1 h on ice. The filtrate was passed through the resin again, drained and the column was washed with 25 mL of wash buffer 1 (40 mM Tris.HCl pH 8.0, 300 mM NaCl and 1% Triton X-100), 25 mL of wash buffer 2 (40 mM Tris.HCl pH 8.0, 300 mM NaCl, 50 mM Na-cholate and 20 mM imidazole), and 25 mL of wash buffer 3 (40 mM Tris.HCl pH 8.0, 300 mM NaCl and 50 mM imidazole). The proteins were eluted from the column with 8 mL of elution buffer (40 mM Tris.HCl pH 8.0, 300 mM NaCl and 400 mM imidazole). The eluate was pipetted into Spectra/Por 3, 3.5 kDa MWCO, 18 mm flat width dialysis membranes and dialyzed against buffer (20 mM Tris.HCl pH 7.4, 100 mM NaCl and 0.5 mM EDTA) overnight.

**Enzyme-coupled ATPase assay**

ATP hydrolytic activity was determined using an NADH enzyme-linked assay (5) adapted for a microplate reader (6), as previously described (3). 50 mL reactions contained assay buffer (TBS for nanodisc-embedded samples; 20 mM Tris. HCl pH 8.0, 150 mM NaCl, 5 mM $MgCl_2$,10% glycerol, 0.05% DDM for detergent-solubilized samples) with 200 mM NADH (Sigma- Aldrich), 20 U/mL lactic dehydrogenase (Sigma-Aldrich), 100 U/mL pyruvate kinase (Sigma-Aldrich), 0.5 mM phosphoenolpyruvate (Alfa Aesar) and different ATP (Sigma-Aldrich) concentrations. The assay were performed at either 37°C or room temperature and fluorescence emission at 340 nm was measured using a SPECTRAmax 250 microplate spectrophotometer equipped with SOFTmax PRO software (Molecular Devices, CA, USA). Readings were taken in ~9 s intervals. The rate of decrease of NADH fluorescence (due to oxidation) was calculated from a linear fit to each 10 min time course and converted to ATP hydrolysis rates with a standard curve obtained using known ADP concentrations. Where indicated, holo and apo MlaC were added in a 5:1 ratio to nanodisc-embedded MlaFEDB complex used. Vanadate was also used at concentrations of 0.6, 1.2 and 2.7 mM. Samples were performed in technical triplicates and data were fit to the built-in Hill equation in OriginPro 2018b.

Both ATP (Sigma-Aldrich) and AMP-PNP (Sigma-Aldrich) were prepared by adjusting the solution pH to pH 8. Sodium orthovanadate (Sigma-Aldrich) was prepared in water adjusted to pH 10. The solution was boiled until clear to ensure presence of vanadate monomers. Upon cooling down to room temperature, the pH was re-adjusted to pH 10. Repeated cycles of boiling and pH adjustment was done until the solution remains clear at pH 10.

**SDS-PAGE, BN-PAGE, immunoblotting and staining**

All samples subjected to SDS-PAGE were mixed with equal amounts of 2X Laemmli reducing buffer. Equal volumes of the unheated samples were loaded onto the gels. Unless otherwise stated, SDS-PAGE was performed according to Laemmli using the using 4-12% Tris.HCl stacking gels (7). After SDS-PAGE, gels were visualized by either Coomassie blue staining (InstantBlue™, expedeon), silver staining (Life Technologies) or subjecting to immunoblotting. BN-PAGE was performed according to published protocols (8) with the usage of 4–20% Tris.HCl polyacrylamide gel. Immunoblotting was performed by transferring protein bands from the gels onto polyvinylidene fluoride (PVDF) membranes (Immun-Blot® 0.2 µm, Bio-Rad) using semi-dry electroblotting system (Trans-Blot®

Turbo$^{TM}$ Transfer System, Bio-Rad). Membranes were blocked by 1X casein blocking buffer (Sigma-Aldrich). α-His antibody (pentahistidine) conjugated to the horseradish peroxidase (HRP) (Qiagen) was used at a dilution of 1:5,000. Rabbit α-MlaC (1) was used at a dilution of 1:500. Donkey α-Rabbit conjugated to HRP (GE Healthcare) was used at a dilution of 1:5000. Luminata Forte Western HRP Substrate (Merck Milipore) was used to develop the membranes and chemiluminescence signals were visualized by G:Box Chemi-XX 6 (Genesys version 1.4.3.0, Syngene). For gels ran with radioactive samples, gels were directly dried via the DryEase® Mini-Gel Drying System (Invitrogen) overnight, after being subjected to either Coomassie blue or silver staining. The dried gels were subsequently imaged by a scanner (CanoScan LiDE 20, Canon).

### Supplementary Figures

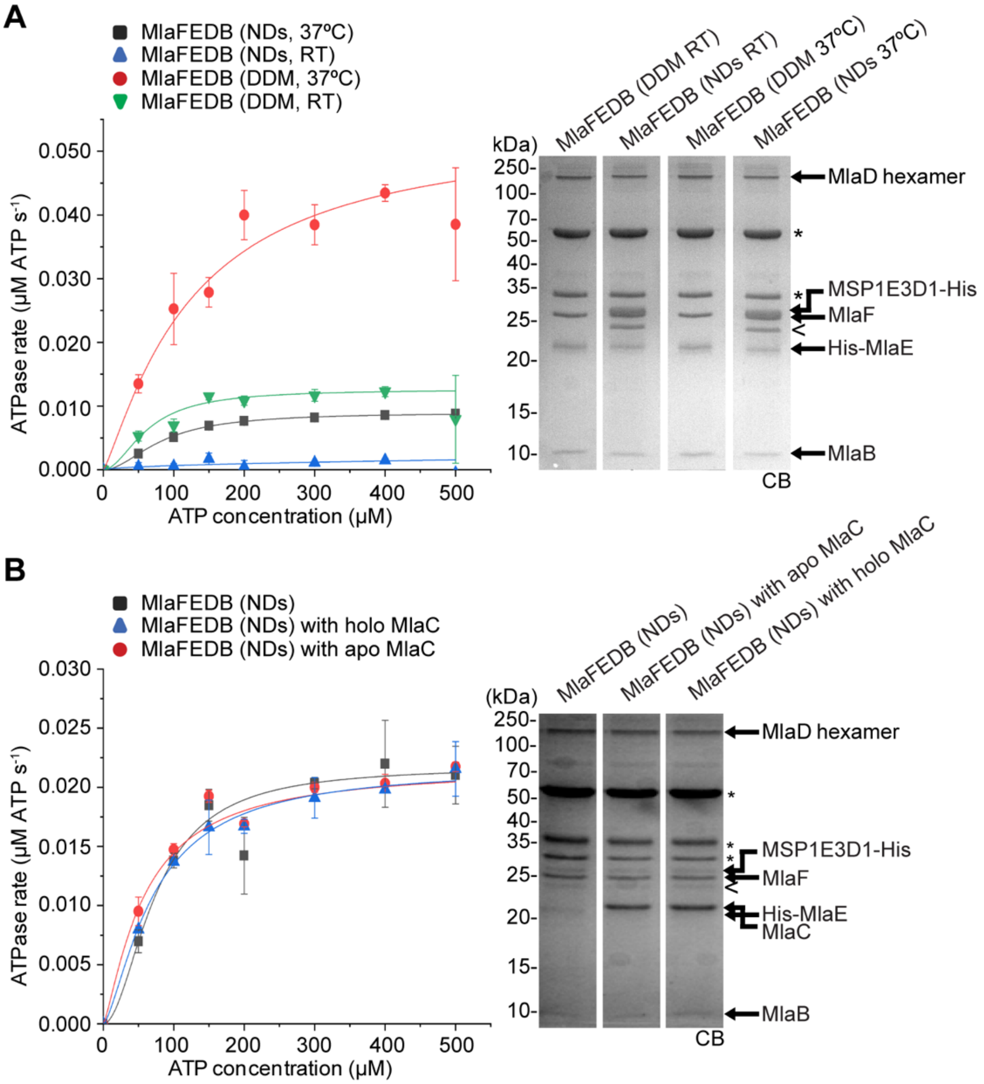


**Figure S1.** Nanodisc-embedded MlaFEDB complex displays high ATP hydrolytic activity at physiological temperature (37°C) but is not further activated by MlaC. (A) Enzyme-coupled ATPase assays of nanodisc-embedded or detergent-solubilized MlaFEDB complexes (0.1 µM) at either 37°C or room temperature. Average ATP hydrolysis rates from triplicate experiments were plotted against ATP concentrations, and fitted to an expanded Michaelis-Menten equation that includes a term for Hill coefficient (n); MlaFEDB (NDs, 37°C) ($k_{cat}$ = 0.090 ± 0.002 µmol ATP $s^{-1}$/µmol complex, $K_m$ = 82.5 ± 4.3 µM, n = 1.9 ± 0.2), MlaFEDB (NDs, RT) ($k_{cat}$ = 0.029 ± 0.024 µmol ATP $s^{-1}$/µmol complex), MlaFEDB (DDM, 37°C) ($k_{cat}$ = 0.540 ± 0.079 µmol ATP $s^{-1}$/µmol complex, $K_m$ = 125.8 ± 40.7 µM, n = 1.2 ± 0.3) and MlaFEDB (DDM, RT) ($k_{cat}$ = 0.126 ± 0.008 µmol ATP $s^{-1}$/µmol complex, $K_m$ = 62.8 ± 7.6 µM, n = 1.9 ± 0.5). (B) Enzyme-coupled ATPase assays of nanodisc-embedded MlaFEDB complexes (0.1 µM) at 37°C in the presence of 5-fold excess of apo or holo MlaC. MlaFEDB (NDs) ($k_{cat}$ = 0.220 ± 0.000 µmol ATP $s^{-1}$/µmol complex, $K_m$ = 75.0 ± 8.0 µM, n = 1.9 ± 0.4), MlaFEDB (NDs with holo MlaC) ($k_{cat}$ = 0.209 ± 0.008 µmol ATP $s^{-1}$/µmol complex, $K_m$ = 67.2 ± 4.0 µM, n = 1.6 ± 0.2), and MlaFEDB (NDs with apo MlaC) ($k_{cat}$ = 0.332 ± 0.207 µmol ATP $s^{-1}$/µmol complex, $K_m$ = 156.8 ± 367.6 µM, n = 0.5 ± 0.4). Errors depicted by bars and ± signs are standard deviations of triplicate data. SDS-PAGE analysis of the complexes used for these assays is shown on the right. *, pyruvate kinase/lactate dehydrogenase enzymes used in coupled assay; <, degraded MSP1E3D1. CB, Coomassie blue staining.

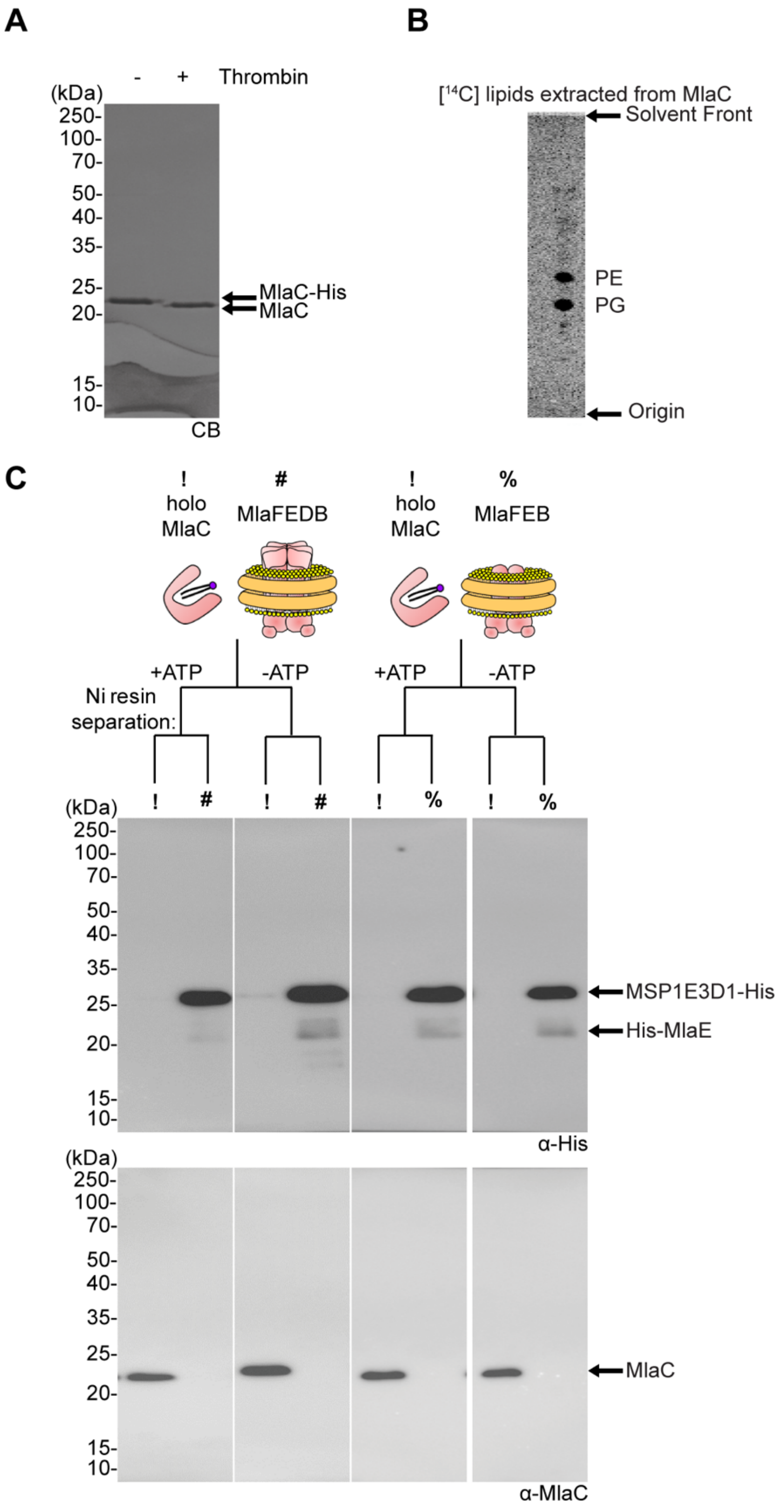


**Figure S2.** Tag-less MlaC is purified bound to [$^{14}$C]-labelled lipids and can be separated cleanly from His-tagged nanodisc-embedded MlaFEDB complex. (A) SDS-PAGE analysis of His-tagged and thrombin-digested tag-less MlaC purified from cells metabolically labelled with [$^{14}$C]-acetate. CB, Coomassie blue staining. (B) Thin-layer chromatography (TLC)/phosphor imaging analysis of the organic extracts from thrombin-digested MlaC, revealing presence of bound [$^{14}$C]-labelled lipids, phosphatidylethanolamine (PE) and phosphatidylglycerol (PG). Lipids are annotated as such because previous work using $^{31}$P NMR analysis indicated that lipids co-purified with MlaC consist of ~50% PG and ~50% PE, with no detectable $^{31}$P signal from cardiolipin (1). (C) Immunoblot analyses using α-His or α-MlaC antibodies, showing clean separation of non-radioactive tag-less holo MlaC (10 μM) and non-radioactive His-tagged nanodisc-embedded complexes (2 μM) by nickel affinity resin. !, holo MlaC; #, nanodisc-embedded MlaFEDB; %, nanodisc-embedded MlaFEB.

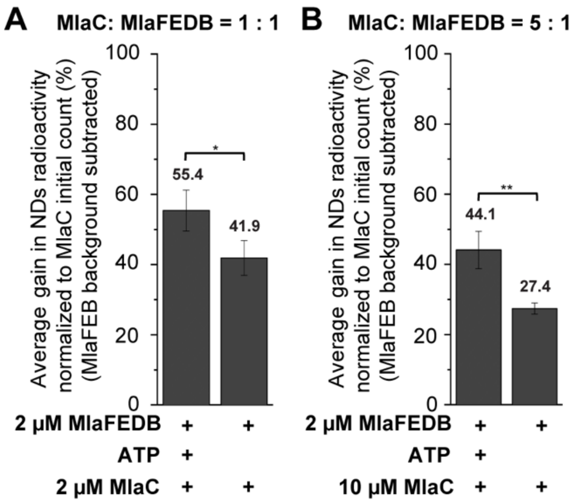


**Figure S3.** Spontaneous transfer of [$^{14}$C]-labelled lipids from MlaC to MlaFEDB decreases when an increased concentration of MlaC is used. Average gains of radioactivity ([$^{14}$C]-lipids) in the indicated nanodiscs from three sets of triplicate experiments, normalized to the initial counts on MlaC when incubated with (A) 2 μM MlaC and (B) 10 μM MlaC. Data from co-incubation of MlaFEB/MlaC are treated as unspecific background transfer/loss and have been subtracted. *, $p < 0.05$; **, $p < 0.01$.

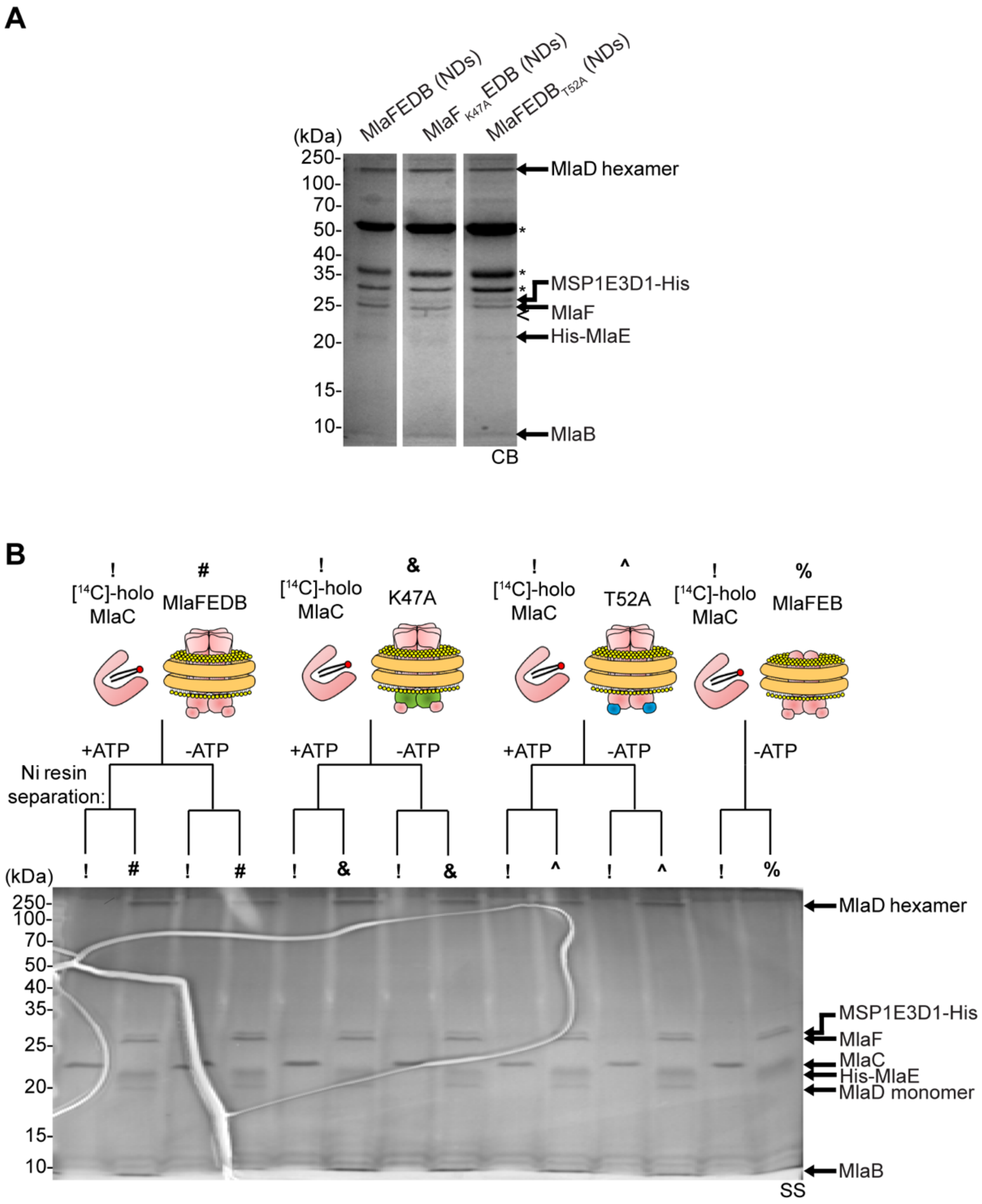
A
MlaFEDB (NDs)
MlaF K47A EDB (NDs)
MlaFEDB T52A (NDs)
(kDa)
250-
100-
70-
50-
40-
35-
25-
20-
15-
10-
MlaD hexamer
*
*
*
MSP1E3D1-His
MlaF
His-MlaE
MlaB
CB
B
!
[14C]-holo
MlaC
#
MlaFEDB
&
K47A
^
T52A
%
MlaFEB
+ATP
-ATP
Ni resin
separation:
(kDa)
MlaD hexamer
MSP1E3D1-His
MlaF
MlaC
His-MlaE
MlaD monomer
MlaB
SS

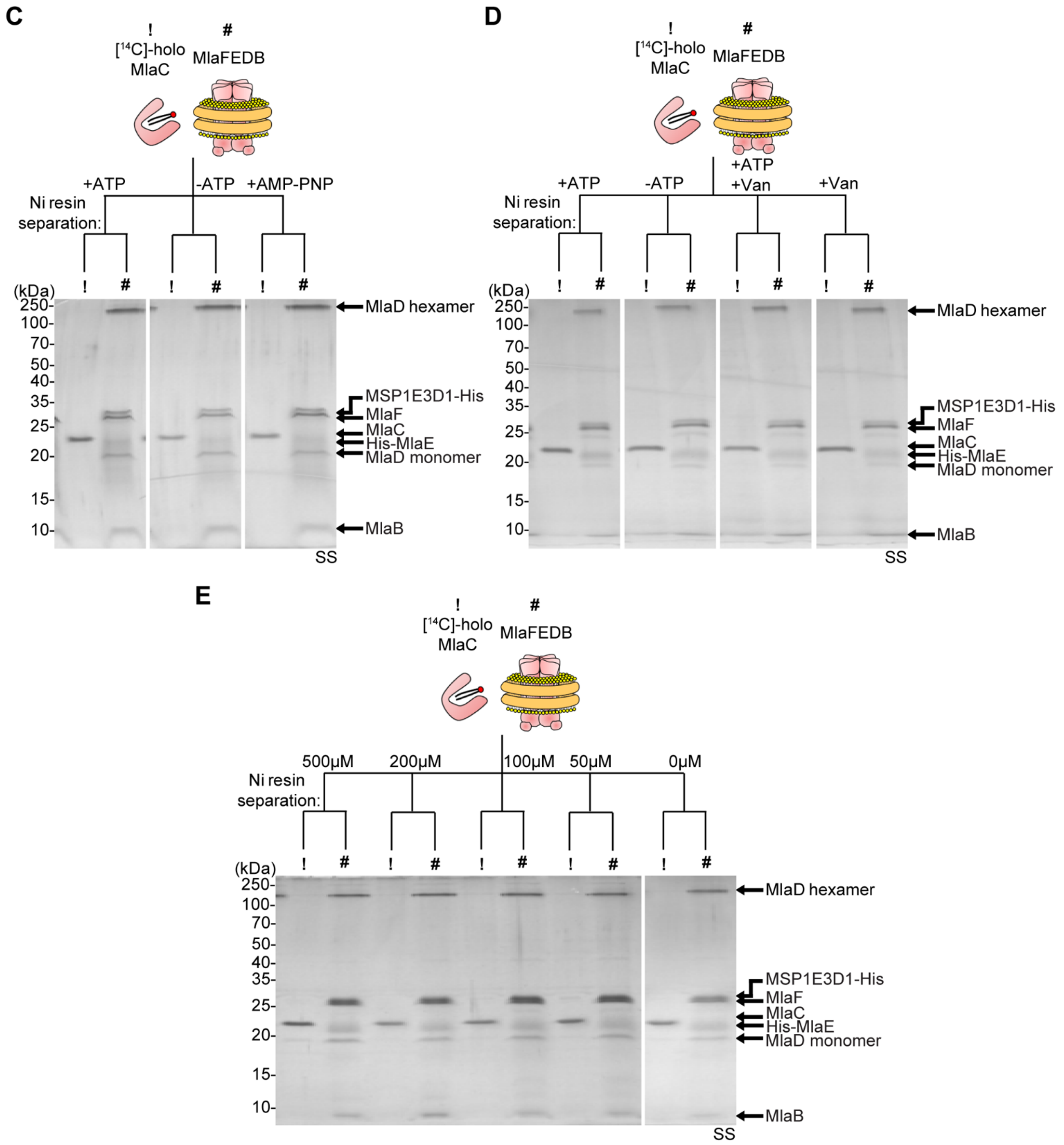


**Figure S4.** Holo MlaC bound with [$^{14}$C]-labelled lipids can be separated from nanodisc-embedded MlaFEDB complexes in assays that probe for the effect of ATP hydrolysis on PL transfer. (A) SDS-PAGE analysis of the complexes used in the enzyme-coupled ATPase assay shown in Figure 3A. *, pyruvate kinase/lactate dehydrogenase enzymes used in coupled assay; <, degraded MSP1E3D1. CB, Coomassie blue staining. (B-E) SDS-PAGE analyses of indicated samples following co-incubation and subsequent nickel affinity-based separation of tag-less [$^{14}$C]-PL bound (holo) MlaC (10 μM) and His-tagged non-radioactive nanodisc-embedded complexes (2 μM), for (B) ATPase mutant complexes, (C) addition of AMP-PNP, (D) addition of vanadate (2.7 mM, and (E) different ATP concentrations as shown in Figure 3B-E. !, [$^{14}$C]-holo MlaC; #, nanodisc-embedded MlaFEDB; &, MlaF$_{K47A}$EDB; ^, MlaFEDB$_{T52A}$; %, nanodisc-embedded MlaFEB; SS, silver staining.

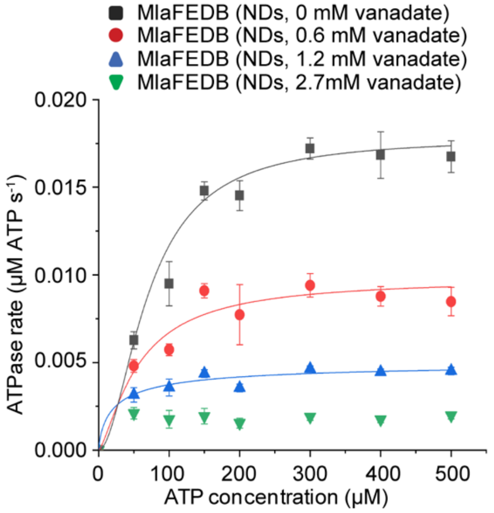


**Figure S5.** Vanadate inhibits the ATPase activity of nanodisc-embedded MlaFEDB complex. Enzyme-coupled ATPase assays of nanodisc-embedded MlaFEDB (0.1 µM) in the presence of 0, 0.6, 1.2 and 2.7 mM vanadate at 37°C. Average ATP hydrolysis rates from triplicate experiments were plotted against ATP concentrations, and fitted to an expanded Michaelis-Menten equation that includes a term for Hill coefficient (n); MlaFEDB NDs (0 mM vanadate) ($k_{cat}$ = 0.178 ± 0.012 µmol ATP $s^{-1}$/µmol complex, $K_m$ = 70.7 ± 9.0 µM, n = 1.8 ± 0.4), MlaFEDB NDs (0.6 mM vanadate) ($k_{cat}$ = 0.098 ± 0.013 µmol ATP $s^{-1}$/µmol complex, $K_m$ = 56.8 ± 12.3 µM, n = 1.4 ± 0.6), MlaFEDB NDs (1.2 mM vanadate) ($k_{cat}$ = 0.051 ± 0.017 µmol ATP $s^{-1}$/µmol complex, $K_m$ = 24.7 ± 20.5 µM, n = 0.7 ± 1.1) and MlaFEDB NDs (2.7 mM vanadate) ($k_{cat}$ = 0.037 ± 0.934 µmol ATP $s^{-1}$/µmol complex). Errors depicted by bars and ± signs are standard deviations of triplicate data. At 2.7 mM vanadate, the ATPase activity of the complex (0.1 µM) is likely fully inhibited; we believe the 'baseline ATPase activity' observed is due to background NADH oxidation induced by vanadate in the enzyme-coupled ATPase assay that we use (9). However, this background is quite low in the context of the high ATP hydrolytic activity observed for the MlaFEDB complex, thus minimally affects the interpretation of vanadate inhibition experiments.

234

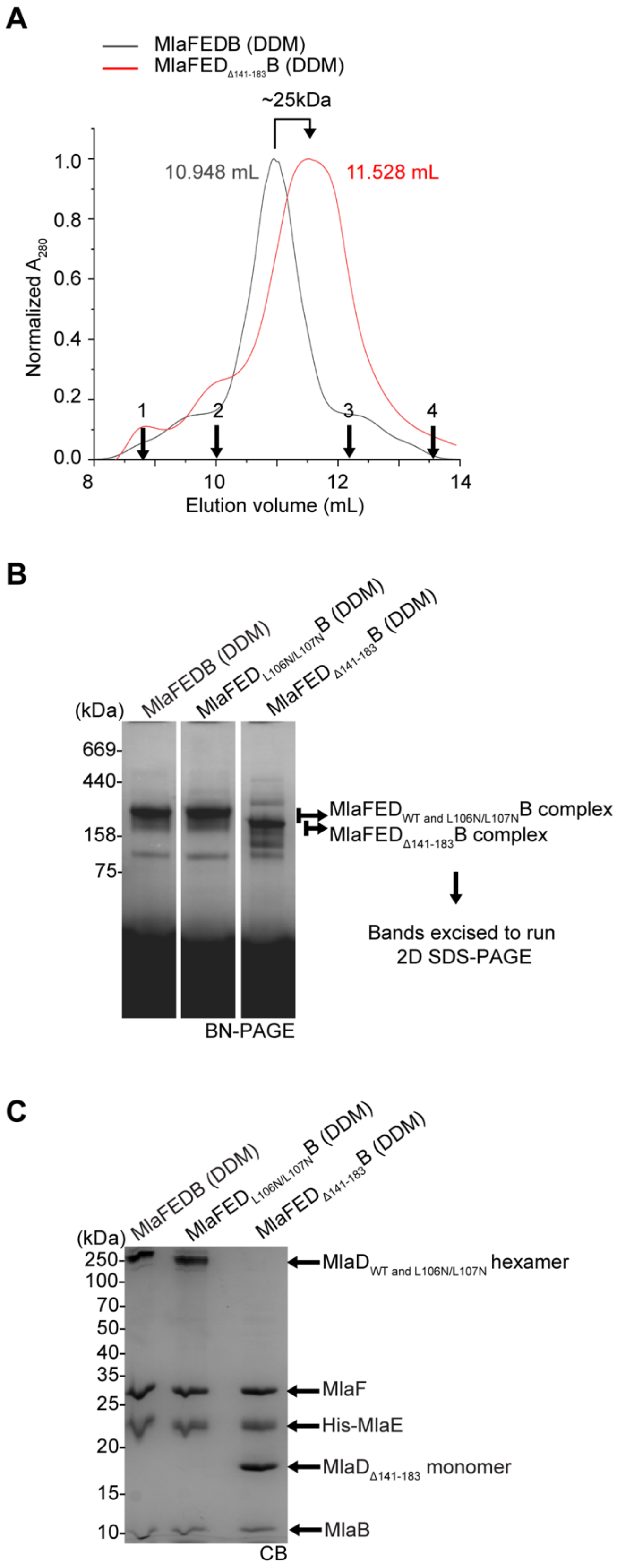
A
MlaFEDB (DDM)
MlaFED Δ141-183 B (DDM)
~25kDa
10.948 mL
11.528 mL
Normalized A280
Elution volume (mL)
1.0
0.8
0.6
0.4
0.2
0.0
8
10
12
14
1
2
3
4
B
(kDa)
MlaFEDB (DDM)
MlaFED L106N/L107N B (DDM)
MlaFED Δ141-183 B (DDM)
669
440
158
75
MlaFED WT and L106N/L107N B complex
MlaFED Δ141-183 B complex
Bands excised to run 2D SDS-PAGE
BN-PAGE
C
(kDa)
MlaFEDB (DDM)
MlaFED L106N/L107N B (DDM)
MlaFED Δ141-183 B (DDM)
250
100
70
50
40
35
25
20
15
10
MlaD WT and L106N/L107N hexamer
MlaF
His-MlaE
MlaD Δ141-183 monomer
MlaB
CB

**Figure S6.** MlaFED$_{\Delta 141\text{-}183}$B forms hexamers in the complex that are non-SDS-resistant, unlike in wildtype MlaFEDB and MlaFED$_{L106N/L107N}$B. (A) Size-exclusion chromatographic (SEC) profiles of detergent-solubilized MlaFEDB (*black*) and MlaFED$_{\Delta 141\text{-}183}$B (*red*) complex. The number beside each peak represents their respective elution volume. The arrows at the axis represent standards (1) thyroglobulin, (2) ferritin, (3) aldolase and (4) conalbumin at their respective elution volumes on a Superdex 200 Increase 10/300 GL column. The difference in elution volumes roughly corresponds to a mass shift of ~25 kDa, consistent with the loss of six C-terminal regions of MlaD, each comprising 43 amino acids (~4.4 kDa). (B) BN-PAGE analysis of SEC-purified MlaFEDB, MlaFED$_{L106N/L107N}$B and MlaFED$_{\Delta 141\text{-}183}$B complex in DDM. The bands indicated were excised and subjected to another SDS-PAGE analysis. (C) SDS-PAGE analysis of the excised bands from (B) shows that MlaFED$_{\Delta 141\text{-}183}$B complex does not form SDS-resistant hexamers unlike that in wildtype and MlaD$_{L106N/L107N}$B. CB, Coomassie blue staining.

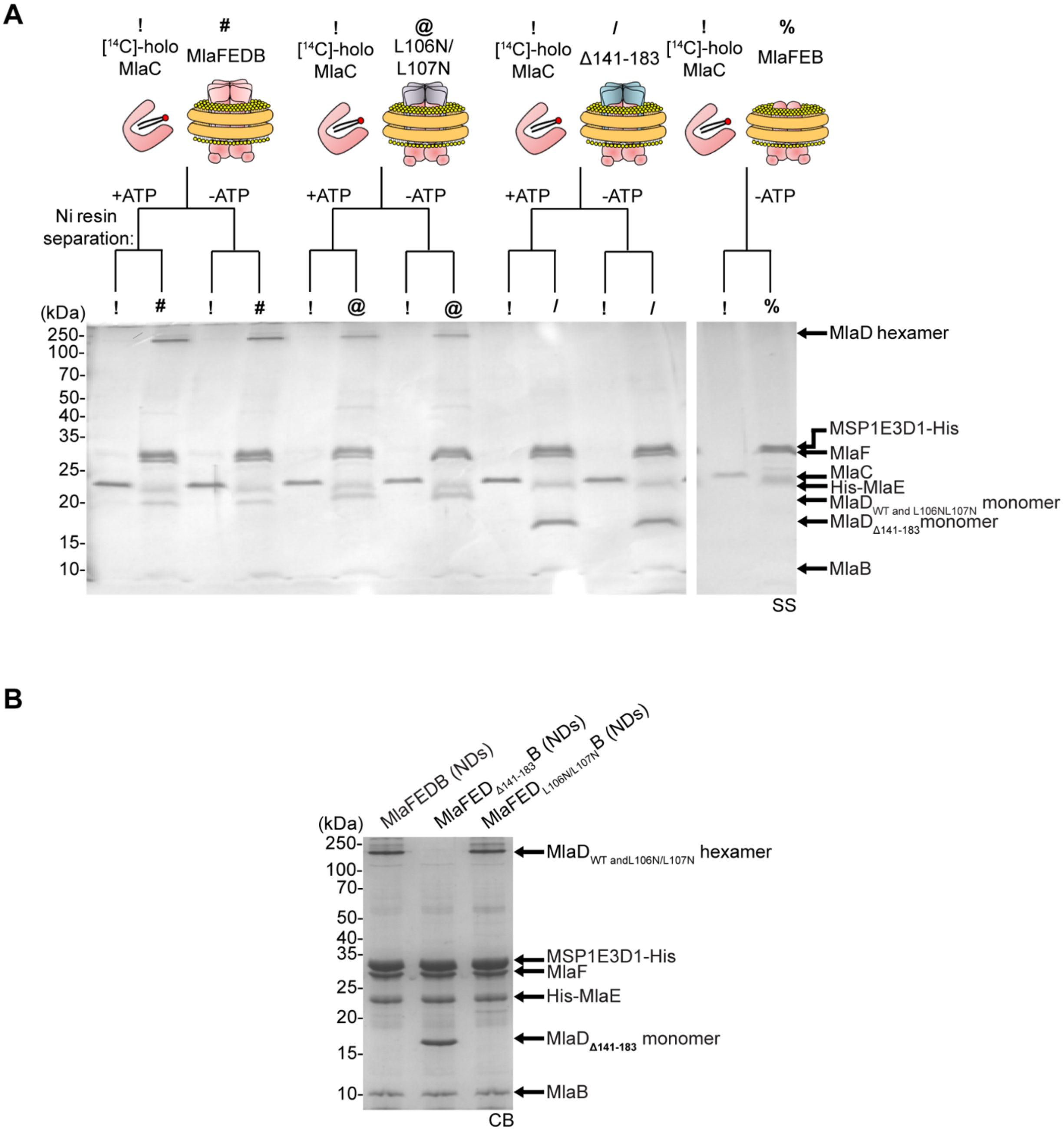


**Figure S7.** Holo MlaC bound with [14C]-labelled lipids can be separated from nanodisc-embedded MlaFEDB, $MlaFED_{L106N/L107N}B$ and $MlaFED_{\Delta141\text{-}183}B$ complexes in assays that probe for the effects of MlaD on PL transfer. (A) SDS-PAGE analysis of indicated samples following co-incubation with or without ATP, and subsequent nickel affinity-based separation of tag-less [14C]-PL bound (holo) MlaC (10 μM) and His-tagged non-radioactive nanodisc-embedded complexes containing MlaD variants (2 μM) as shown in Figure 4A. !, [14C]-holo MlaC; #, nanodisc-embedded MlaFEDB; @, $MlaFED_{L106N/L107N}B$; /, $MlaFED_{\Delta141\text{-}183}B$; %, nanodisc-embedded MlaFEB; SS, silver staining. (B) SDS-PAGE analysis of the complexes used in the enzyme-coupled ATPase assay shown in Figure 4B. *, pyruvate kinase/lactate dehydrogenase enzymes used in coupled assay. CB, Coomassie blue staining.

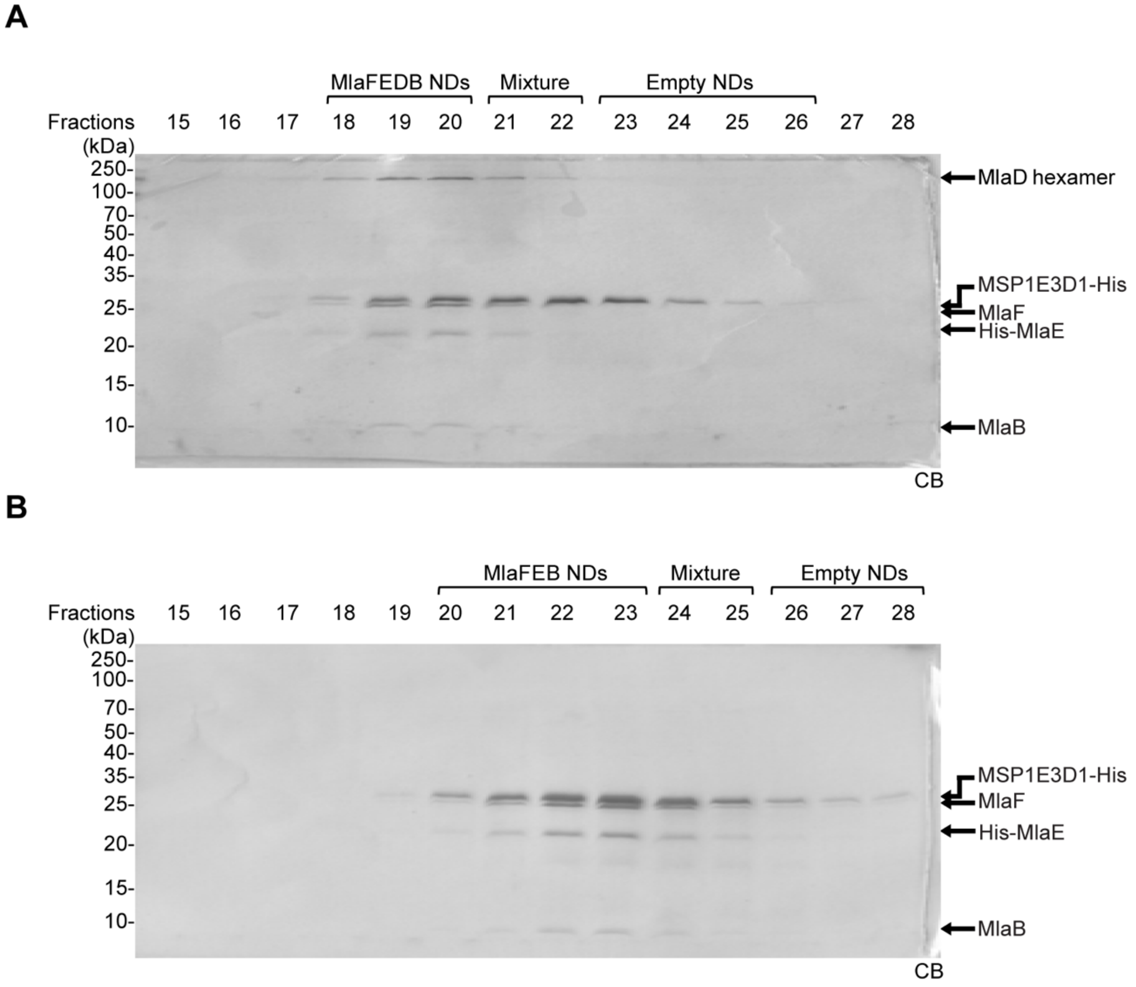


**Figure S8.** MlaFEDB and MlaFEB complexes are reconstituted into nanodiscs containing [$^{14}$C]-labelled PLs and purified by size exclusion chromatography. SDS-PAGE analyses of samples from indicated collected fractions (0.5 mL each) of reconstituted radioactive nanodisc-embedded (A) MlaFEDB complex and (B) MlaFEB complex, following purification on a self-packed Superose 6 column. CB, Coomassie blue staining.
